# Rostrocaudal patterning and neural crest differentiation of human pre-neural spinal cord progenitors in vitro

**DOI:** 10.1101/2020.06.16.155564

**Authors:** Fay Cooper, George E Gentsch, Richard Mitter, Camille Bouissou, Lyn Healy, Ana Hernandez Rodriguez, James C Smith, Andreia S Bernardo

## Abstract

The spinal cord emerges from a niche of neuromesodermal progenitors (NMPs) formed and maintained by Wnt/FGF signals at the posterior end of the embryo. NMPs can be generated from human pluripotent stem cells and hold promise for spinal cord replacement therapies. However, NMPs are transient, which complicates the full range production of rostrocaudal spinal cord identities *in vitro*. Here we report the generation of NMP-derived pre-neural progenitors (PNPs) with stem cell-like self-renewal capacity. PNPs maintain pre-spinal cord identity by co-expressing the transcription factors SOX2 and CDX2, and lose mesodermal potential by downregulating TBXT. For 7 to 10 passages PNPs divide to self-renew and to make trunk neural crest (NC), while gradually adopting a more posterior identity by activating colinear *HOX* gene expression. This HOX clock can be halted at the thoracic level for up to 30 passages by blocking the trunk-to-tail transition through GDF11-mediated signal inhibition.

## INTRODUCTION

Pluripotent stem cells (PSCs) have become an important tool for the study of mammalian development. Directed differentiation of PSCs *in vitro* has given significant insights to the signals and gene regulatory networks which are important for cell fate decisions (Baillie-Benson et al., 2020). In particular, PSC-derived neural stem cells (NSCs) are often an effective starting point in understanding both neural development and disease, and have great potential for use in regenerative medicine (Snyder, 2017). However, to use *in vitro* NSCs in this way, differenation protocols must recapitulate *in vivo* by following the correct developmental route and must reproducibly generate a well charactered NSC population. The therapeutic value of human NCSs hinges on them adopting the correct region-specific identities and adapting properly to their local microenvironments (Kadoya et al., 2016, Kumamaru et al., 2018, Nagoshi et al., 2019). For example, patients with motor neuron disease or spinal cord injuries often display lesions in specific neuronal cell types. Hence, effective therapeutic repair depends on developing protocols that reliably generate these neuronal subtypes from induced pluripotent stem cells (iPSCs)(Trawczynski et al., 2019, Nijssen et al., 2017).

Not surprisingly, the development of parts of our central nervous system *in vitro* has been inspired by our knowledge of mammalian neurogenesis. Forebrain and midbrain develop from the anterior neural plate, a naïve tissue neuralised by the underlying axial mesoderm through the release of TGF-β inhibitors (Cajal et al., 2012, Mathis and Nicolas, 2000). Spinal cord arises from a progenitor pool of neuromesodermal progenitors (NMPs) that reside in the caudal lateral epiblast/node streak border and later the chordoneural hinge (Wilson et al., 2009). NMPs are bi-potent and give rise to both the posterior neural tube and adjacent somite-forming paraxial mesoderm (Cambray and Wilson, 2002, Cambray and Wilson, 2007, Delfino-Machin et al., 2005, Tzouanacou et al., 2009, Brown and Storey, 2000). NMPs are maintained by the synergistic action of FGF and Wnt signals which activate the co-expression of the transcription factors TBXT, SOX2 and CDX2. TBXT and SOX2 are mutually antagonistic cell fate determinants for the mesodermal and neuroectodermal germ layers, respectively (Wymeersch et al., 2016, Henrique et al., 2015, Tsakiridis et al., 2014, Gouti et al., 2017, Koch et al., 2017). CDX2 conveys increasingly more posterior identity to NMP descendants by inducing colinear *HOX(1-13)* gene expression during axial elongation (van den Akker et al., 2002, van de Ven et al., 2011, Neijts et al., 2017, Amin et al., 2016). The human HOX genes are expressed in a spatial and temporal order that is colinear with their physical 3’ to 5’ genomic position, and assign overlapping regional identity to the brain and vertebral segments of the spinal cord: HOX1-5, hindbrain; HOX4-6, cervical; HOX6-9, thoracic and HOX10-13, lumbosacral (Philippidou and Dasen, 2013).

As the rostrocaudal axis elongates, NMPs that enter the primitive streak downregulate SOX2, upregulate TBX6, and contribute to the developing somites (Takemoto et al., 2011, Javali et al., 2017). Their alternative commitment to pre-neural progenitors (PNPs) begins in the pre-neural tube (PNT), located immediately rostral to the NMP niche (Diez del Corral et al., 2002). In the PNT, cells no longer express *TBXT*, but maintain expression of *SOX2* and *NKX1-2* (Olivera-Martinez and Storey, 2007, Storey et al., 1998). Neurogenic genes such as *PAX6* and *NEUROG2* are not robustly expressed yet in this region due to the repressive effect of continued FGF signalling on retinoic acid (RA) production (Lunn et al., 2007, Diez del Corral et al., 2003). The next step of neural commitment is then prompted by the exposure of PNPs to RA from the adjacent somites as they migrate out of the PNT region and into the neural tube. The switch from FGF to RA-mediated signalling alleviates repression of the neural transcription factors *PAX6* and *IRX3* and down regulates *NKX1-2* (Sasai et al., 2014, Diez del Corral et al., 2003, Shum et al., 1999).

Attempts have been made *in vitro* to recapitulate the developmental pathways leading to anterior or posterior NSCs. Brain forming anterior NSCs can be generated from human PSCs (hPSCs) via dual TGF-β (Activin/BMP) inhibition (Chambers et al., 2009). Initial attempts to generate spinal cord progenitors relied on posteriorising anterior NSCs through exposure to retinoic acid (Mazzoni et al., 2013, Wichterle et al., 2002, Lee et al., 2007, Li et al., 2005). This yielded neural derivatives as far posterior as hindbrain and upper cervical regions, primarily through saltatory expression of *HOX(1-5)* genes. Following these studies, and consistent with *in vivo* evidence, combined Wnt and FGF stimulation efficiently converted mouse and human PSCs into NMP-like cells (Turner et al., 2014, Gouti et al., 2014, Lippmann et al., 2015, Frith et al., 2018, Verrier et al., 2018, Peljto et al., 2010). Neural progenitors derived from NMP-like cells and generated using Wnt/FGF stimulation are capable of undergoing a more complete range of regionalisation along the rostrocaudal axis, generating neural progenitors up to lumbar identity (HOX10) (Lippmann et al., 2015, Kumamaru et al., 2018, Wind et al., 2020). Furthermore, NMP-like cells have become very informative in studying the intricate cell fate decisions and dynamics of spinal cord formation (Metzis et al., 2018, Gouti et al., 2017, Gouti et al., 2014, Edri et al., 2019, Rayon et al., 2020).

Work using hPSCs to study spinal cord formation is still preliminary. Here we describe the *in vitro* conditions which commit hPSC-derived NMPs to PNPs. These PNPs are stable for up to 10 passages (30 days). They can acquire the full range of rostrocaudal identities, including the most posterior (sacral) identity represented by HOX11-13 gene expression, produce trunk neural crest (NC) and region-specific spinal cord tissue (e.g. motor neurons and interneuron subtypes). Interestingly, the culture of thoracic PNPs can be massively extended by suppressing TGF-β/GDF11-mediated signalling, which in line with previous *in vivo* findings blocks the trunk-to-tail transition (Aires et al., 2019, Jurberg et al., 2013). Together, we present a well characterised, reproducible and simple protocol which holds the potential to model several aspects spinal cord and trunk NC formation *in vitro*.

## RESULTS

### Optimising the generation of NMP-like cells from hPSCs through Wnt modulation

Previous studies have shown that Wnt/FGF signalling causes mouse and hPSCs to adopt neuromesodermal bipotency (Turner et al., 2014, Gouti et al., 2014, Lippmann et al., 2015, Frith et al., 2018, Verrier et al., 2018). Human NMP protocols differ in both the magnitude and time window of Wnt stimulation, as well as with respect to the addition of other signal modulators including FGF (Figure S1A)(Wang et al., 2019, Gouti et al., 2014, Frith et al., 2018, Edri et al., 2019, Verrier et al., 2018, Lippmann et al., 2015, Gomez et al., 2019, Kumamaru et al., 2018, Denham et al., 2015). To find the critical Wnt signalling threshold for the generation of NMP-like cells from the WA09 (H9) hESC line, cells were seeded at a fixed density and 24h later exposed to increasing concentrations of the canonical Wnt agonist CHIR99021 (CHIR) while keeping FGF2 ligands constant (Figure 1A). Our culture medium lacked the retinoic acid (RA) precursor vitamin A (retinol) and contained the pan-RA receptor (RAR) inverse agonist AGN193109 (AGN) (Klein et al., 1996). RA neuralises multipotent cells, so its degradation by CYP26A1 is essential for NMP maintenance (Sakai et al., 2001, Abu-Abed et al., 2001, Martin and Kimelman, 2010). Yet, the RA receptor gamma (RARγ) is highly expressed in NMPs suggesting that transcriptional repression mediated by RARγ in the absence of its ligand supports NMPs and rostrocaudal axis elongation (Janesick et al., 2014). AGN addition reduced aldehyde dehydrogenase (ALDH) activity indicating that endogenous RA synthesis was decreased (Figure S1B).

**Figure 1:**
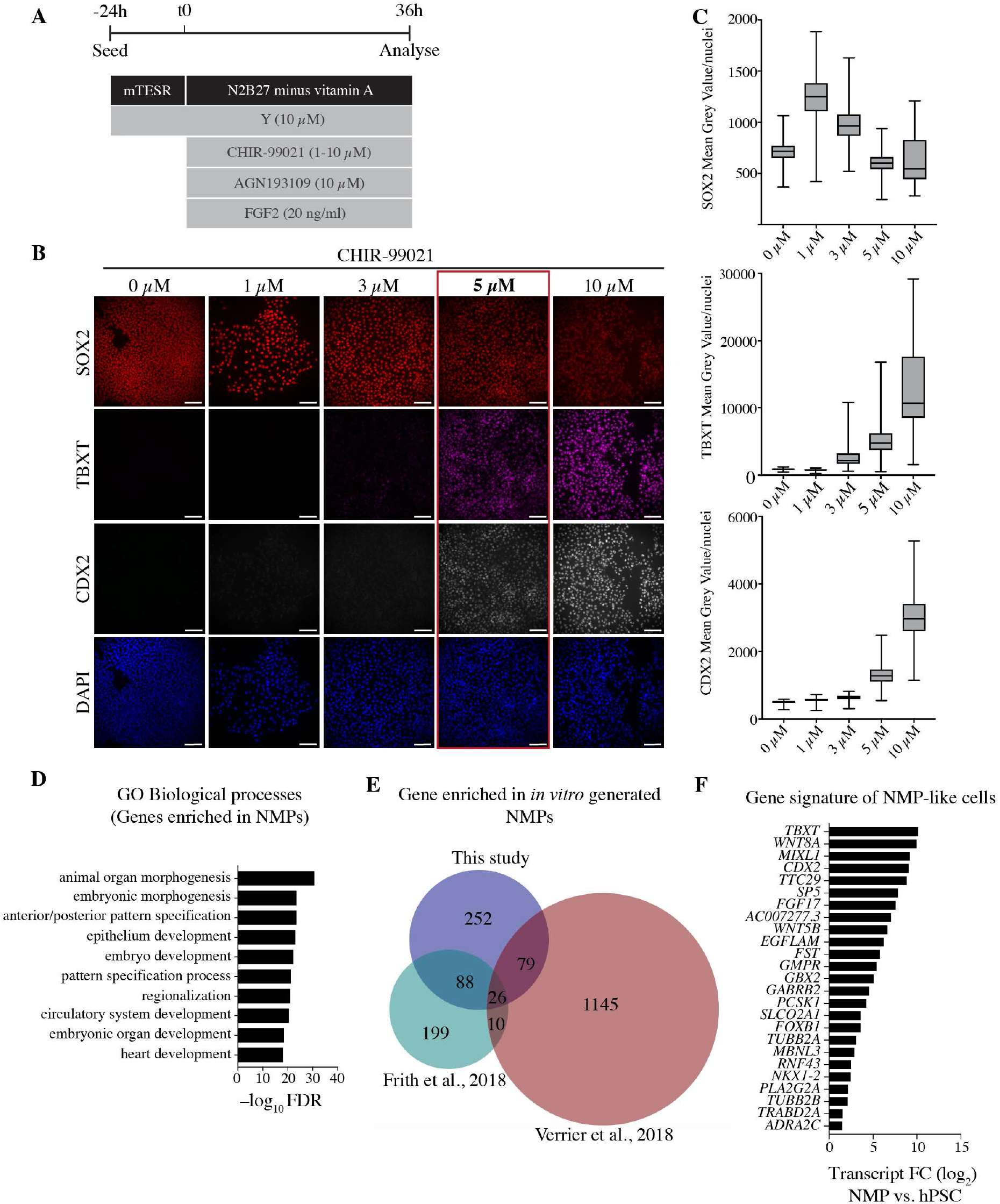
NMP-like cells are induced by intermediate Wnt signalling in the presence of FGF and inhibited RA signalling. (A) Tissue culture scheme for optimising NMP generation from hPSCs. hPSCs are plated 24h before exposure to FGF2 (20ng/ml), CHIR-99021 (0-10 μM), AGN193109 (10 μM) and Y-27632 (10 μM) for 36h. B) Representative immunostaining of 36h cultures treated as shown in (A), showing characteristic NMP markers SOX2 (red), TBXT (magenta), CDX2 (grey) and the nuclear stain DAPI (blue) under different CHIR-99021 concentrations. Scale bars, 100 μm. C) Box-plots showing mean grey value/nuclei quantified from repeat experiments as shown in (B). Each plot show data points collected from 2-4 experiments (>200 nuclei). D) Biological process GO analysis for genes significantly upregulated in NMPs compared to pluripotent hESCs. The top 10 biological process terms with the corresponding Benjamini and Hochberg adjusted p-values (FDR) are shown. E) Venn diagram showing the overlap of significantly upregulated genes in NMPs as reported in this study, Frith et al., (2018) and Verrier et al., (2018). F) Graph showing transcriptional fold change (FC) within the dataset of this study, of 26 genes commonly upregulated in NMPs according to Venn diagram in (E).

After 36h, when cultures reached confluency, cells were analysed for SOX2, TBXT and CDX2 expression by immunofluorescence (Figure 1B,C). Low concentrations of CHIR (0-1μM) caused cells to express high SOX2 and to be negative for TBXT and CDX2. At 3μM CHIR, TBXT and CDX2 protein became detectable in some cells. At 5-10μM CHIR, TBXT and CDX2 levels were further elevated, while SOX2 expression decreased with increasing concentrations of CHIR. Bearing in mind the role of POU5F1 (also known as OCT4) in maintaining pluripotency and axis elongation (Aires et al., 2016, Gouti et al., 2017), we also analysed expression of this protein at increasing CHIR concentrations. As expected, when cells were treated with rising CHIR concentrations, OCT4 expression was lost (Figure S1C,D). Based on the co-expression of OCT4, SOX2, CDX2 and TBXT proteins, we determined that 5μM CHIR was the optimal concentration to generate NMP-like cells from H9 hESCs at this cell density in 36h. We could also reliably generate NMP-like cells from WA01 (H1) hESCs and the AICS-ZO1-GFP iPSC line, which also required intermediate (but different) levels of Wnt activation (Figure S2A,B,E,F). These data show that optimising the magnitude of Wnt signalling is important for obtaining NMP-like cells from different PSC lines.

### Transcriptional profiling reveals a common NMP gene set

To further characterise our NMP-like cells, Wnt/FGF-induced transcriptional changes in H9 hPSCs were quantified by bulk RNA sequencing (RNA-Seq). 1,367 genes were significantly differentially expressed between hESC and NMP stages (445 up and 922 down; FDR <1%, a fold change of at least ± 2, and a base mean >100) (Supplementary file 1). The biological processes most significantly enriched within upregulated genes included ‘anterior-posterior pattern specification’ and ‘regionalisation’, processes which reflect the roles of NMPs *in vivo* (Figure 1D). To define a common gene set expressed by *in vitro* NMPs, we compared our gene list of upregulated genes with two other NMP-related gene expression studies (Verrier et al., 2018, Frith et al., 2018). The comparison revealed 26 genes that were consistently upregulated in all three studies (Figure 1E, F). Among these were well-established NMP markers such as TBXT, WNT8A, CDX2, FGF17, FST and NKX1-2 (Figure 1F). Several novel genes were also identified, including AC007277.3, a long non-coding transcript, and TTC29 and EGFLAM, all of which may be useful as NMP markers. Overall, our results show that hPSC-derived NMPs generated in the absence of RA signalling express known *in vivo* NMP marker genes and share a distinct gene signature with other *in vitro* hPSC-derived NMPs.

### Prolonged culture of NMPs results in loss of mesodermal potency and the emergence of epithelial SOX2^+^/CDX2^+^ colonies

NMPs have previously been maintained in culture for up to seven days (Lippmann et al., 2015), but it is necessary to culture them for longer than this to create enough cells for developmental and therapeutic assays. We sought to extend the culture of spinal cord progenitors by generating the posterior (SOX2^+^/CDX2^+^) equivalent of anterior (SOX2^+^/OTX2^+^) NSCs. To this end we dissociated and replated NMP-like cells at low density at 36h, supressed RA signalling (by removal of vitamin A from the medium and treatment with AGN) and continued Wnt/FGF treatment to minimise mesodermal commitment while halting early neural commitment (Figure 2A).

**Figure 2:**
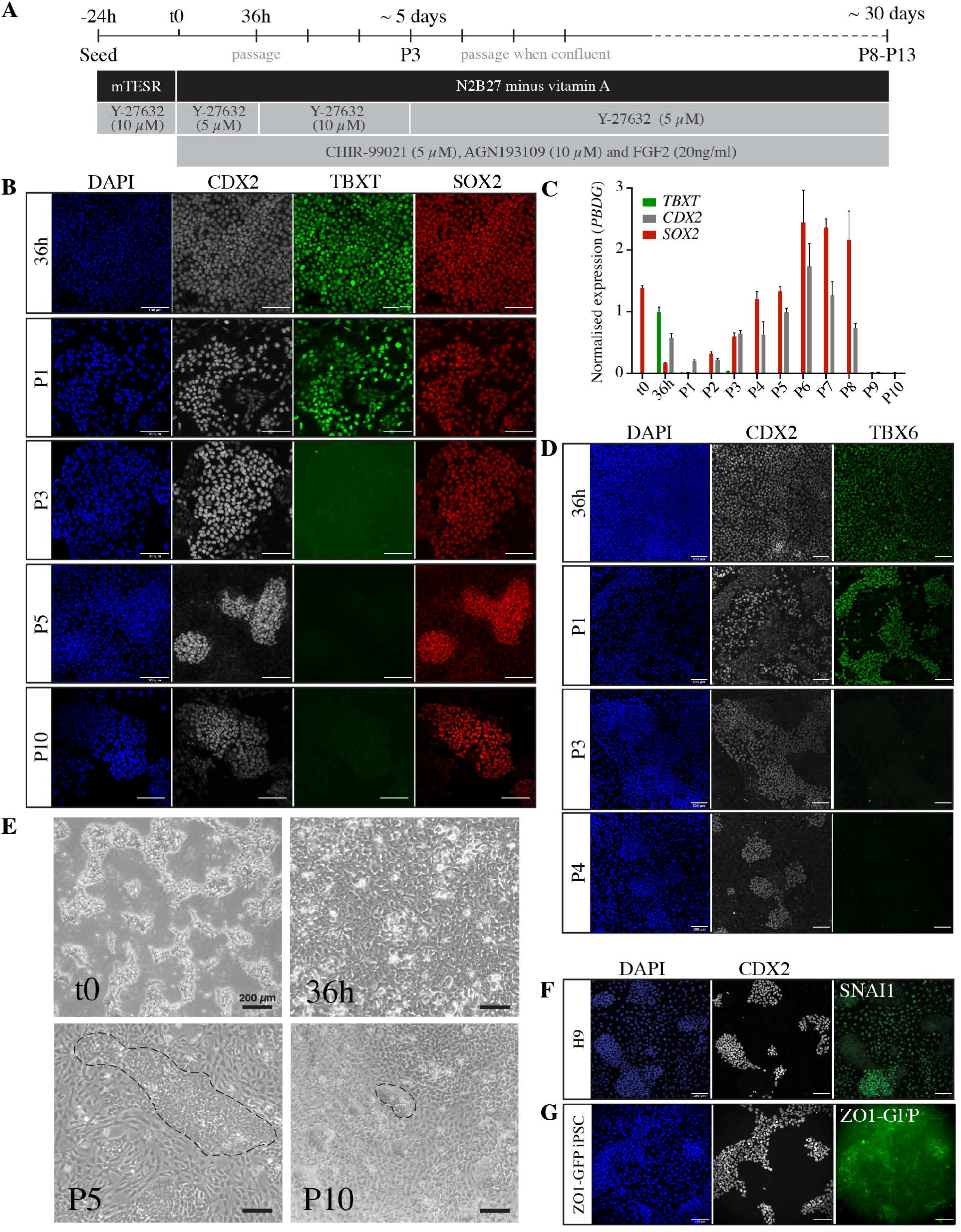
Long term culture of NMPs in the presence of Wnt/FGF and inhibited RA signalling generates epithelial SOX2^+^/CDX2^+^ cell colonies. A) Tissue culture scheme for generating NMPs and maintaining neural progenitors *in vitro*. Cells are passaged at 36h and subsequently passaged at 80-90% confluency for up to 13 passages in FGF2 (20ng/ml), CHIR-99021 (5 μM), AGN193109 (10 μM) and Y-27632 (10 or 5 μM). B) Representative immunostaining of CDX2 (grey), TBXT (magenta), SOX2 (red) and nuclear stain DAPI (blue) at increasing stages of tissue culture (36h, passage (P)1, P3, P5 and P10). Scale bars, 100 μm. C) Transcriptional analysis (RT-qPCR) of NMP markers at each passage up to passage 10. Expression levels are normalised to the reference gene *PBDG*. Error bars show SD, (n=3 technical replicates). Data are *representative* of three independent *experiments*, biological replicates provided in Figure S3A,B. D) Representative immunostaining of TBX6 (green), CDX2 (grey) and nuclear stain DAPI (blue) at 36h, P1 and P3. Scale bar, 100 μm. E) Representative brightfield images of cells at the indicated stages. Dashed lines in P5 and P10 outline examples of a compact epithelial colonies, which are surrounded by flat mesenchymal cells. Scale bar, 200 μm. F) Representative immunostaining of CDX2 (grey), SNAI1 (green) and the nuclear stain DAPI (blue) at passage 5. Scale bar, 100 μm. G) Representative immunostaining of CDX2 (grey), GFP (ZO1-mEGFP iPSC, green) and the nuclear stain DAPI (blue) at passage 5. Scale bar, 100 μm.

Using immunofluorescence and RT-qPCR, we showed that these culture conditions maintain a SOX2^+^/CDX2^+^ cell population between up to 10 passages, corresponding to ~30 days (Figure 2B,C and S3A, B). While *SOX2* and *CDX2* transcripts were detected, their levels varied between experiments and normally dropped between P7 and P10 (Figures 2B,C and S3A,B). Similar observations were made when using H1 hESC and AICS ZO1-mEGFP iPSCs (Figure S2C,G). After one passage (P1) the cultures were heterogeneous with some cells expressing the NMP-characteristic TBXT^+^/SOX2^+^/CDX2^+^ signature. By P3, TBXT and its immediate downstream target TBX6 were undetectable, but most cells continued to express CDX2 and SOX2, suggesting a loss of mesodermal and a maintenance of neural potency (Figure 2B-D).

By P5, the cell population had segregated into two types, as judged by bright-field and immunofluorescence imaging (Figure 2B and 2E): one formed compact SOX2^+^/CDX2^+^ cell colonies, while the other was negative for SOX2/CDX2 and had acquired mesenchymal characteristics such as cell spreading and SNAI1 expression (Figure 2F). The SOX2^+^/CDX2^+^ cells appeared to be epithelial, based on the accumulation of mEGFP-tagged zona occludens (ZO)-1 at tight junctions in transgenic AICS iPSCs (Figure 2G). Together, our results showed that persistent Wnt/FGF signalling without RA converts hPSCs via the transient NMP state into a semi-stable epithelial SOX2^+^/CDX2^+^ cell colonies that could be maintained for 7-10 passages.

### NMPs form neural progenitors and NC derivatives over time

To investigate gene expression changes during the transition of NMP-like cells into epithelial and mesenchymal populations, we profiled the transcriptomes of our cultures by bulk RNA-Seq across twelve time points from 24h after seeding hESCs (time 0, t0) to P10. Analysis of principal components 1 and 2 (PC1 and PC2) showed that most biological replicates (n=2-3) clustered together and PC1 (43% variation) separated according to the duration between time points (Figure S4A). Some outliers were identified, which we presume to be a reflection of biological variation in our experiments. In support of this, outliers such P1.r1 and P2.r1 associated with the previous passage, such that P2.r1 clustered more closely to P1.r2 and P1.r3, suggesting that replicate 1 (r1) differentiated through the same transitions, but at a slower pace than r2 and r3.

Next, k-means hierarchical clustering was applied to all gene-specific profiles that were significantly different over at least two consecutive time points. Each of the gene clusters showed a distinct transcriptional behaviour over time (Figure 3A, Supplementary file 2). The genes of each cluster were analysed for enriched gene associated biological processes in the form of gene ontology (GO) terms and the most significant four GO terms are listed in Figure 3B (Supplementary file 3). Clusters 2 (C2) and 6 (C6) showed elevated gene expression from P1 to P8, when cells robustly expressed SOX2 and CDX2. Consistent with the role of CDX2 in regulating colinear *HOX* gene expression, *CDX2* and *HOX(1-* genes were grouped together in C2, which showed ‘regionalization’ as the most enriched biological process (Neijts et al., 2017, Amin et al., 2016). Conversely, *SOX2* was clustered with other neural fate determinants including *SOX21, SP8* and *GBX2* in C6 and thus, this cluster was linked strongly with various biological functions of neurogenesis. (Luu et al., 2011, Sandberg et al., 2005, Li et al., 2014). As expected, the most posterior HOX genes were found in C4 and C9, which showed a peak of expression around P7-P8 and P9-P10, respectively. This was in line with previous findings indicating *HOX13* genes retro-inhibit anterior *HOX* and *CDX2* transcription (Denans et al., 2015). Thus, we observed full colinear *HOX(1-13)* gene expression across ten passages (Figures 3C,D). The onset of terminal *HOX* gene expression varied in later passages, possibly reflecting slight variation in differentiation rates between experiments (Figure 3A,C, Figure S5A,B). A similar collinear HOX gene expression pattern was noted when using H1 hESC and AICS ZO1-mEGFP iPSCs (Figure S2D,H). In parallel with the onset of terminal HOX expression, C4 and C9 included genes with elevated expression at P9 and P10 (Figure 3A). These clusters were enriched for differentiated tissues such as the skeletal system (C9) and the circulatory system (C4) suggesting that cells at P7/P8 start to differentiate and this provides a genetic explanation for the decrease in cell viability and the increase in cell spreading at late passages (Figure 3A,B,E).

**Figure 3:**
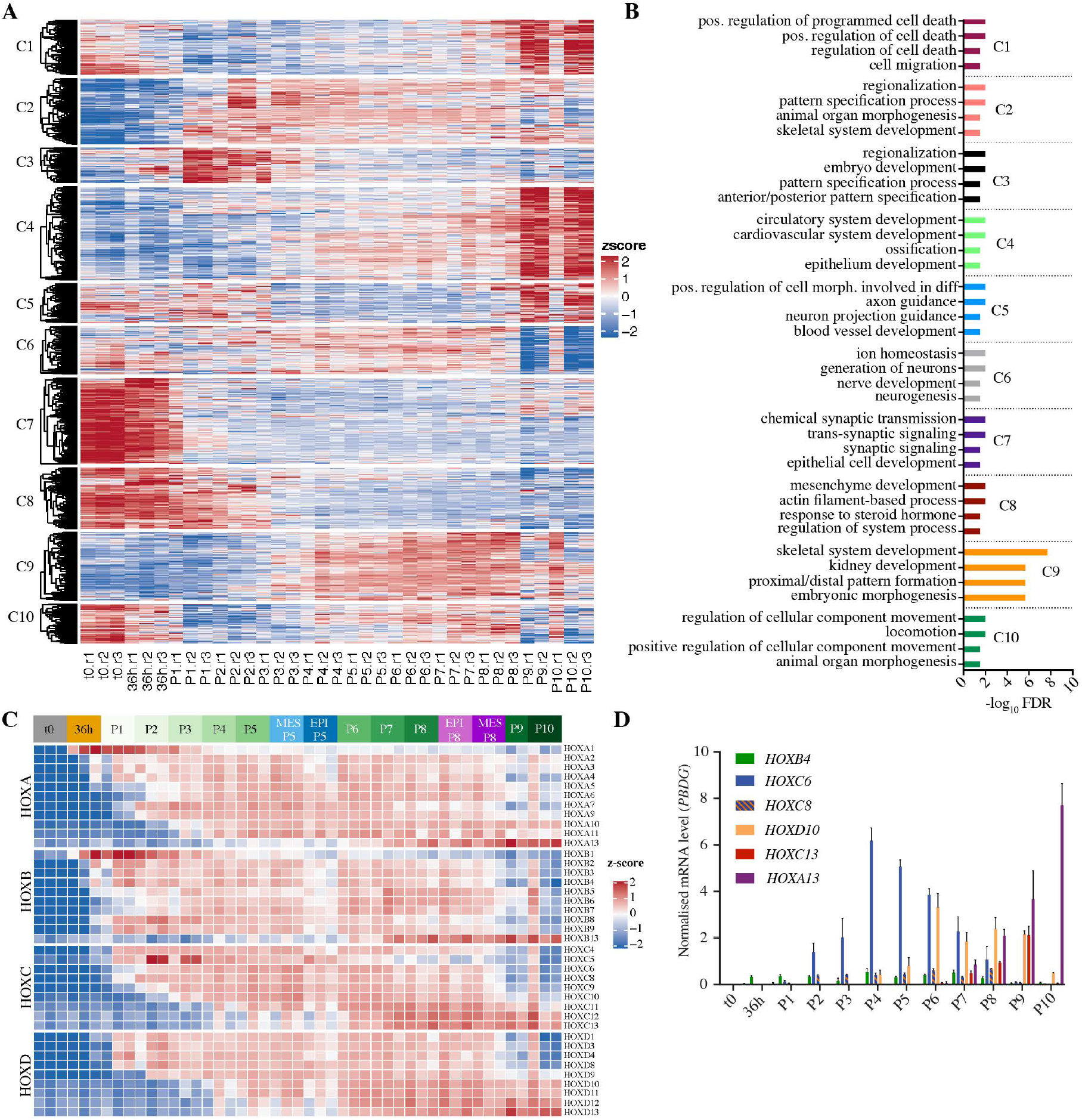
RNA-Seq analysis indicates NMPs transition to neural progenitors and NC derivatives. A) Heatmap showing dynamically expressed genes (z-score) sorted into 10 clusters (C1-10) using k-means hierarchical clustering. Each cluster represents a different temporal expression pattern. B) Biological processes GO analysis for gene sets in each cluster shown in (A). The corresponding Benjamini and Hochberg adjusted p-values (FDR) are shown. C) Heatmap of expressed *HOX(A-D)* genes (z-score) across each time point including enriched epithelial (EPI) and mesenchymal (MES) samples at P5 and P8. D) Transcript levels of selected *HOX* genes as measured by RT-qPCR. Expression level was normalised to the reference gene *PBGD*. Error bars show SD, (n=3 technical replicates). Data are representative of three independent experiments, replicates provided in Figure S5A,B.

Similar to C4, C1 consisted of genes upregulated at P9/P10. C1 and C4 genes were enriched for cell death, cell migration and NC-related biological processes such as ossification, suggesting some loss of cell viability and the onset of cell differentiation in these later passages (Figure 3A,B). Together, these results suggest that cells become NC-like and then terminally differentiate, which would be in keeping with the crest-related tissue types identified within the GO term analysis of C4 and C9. This is also consistent with the decrease in cell viability, which we observe towards passage 10. NMP-like cells appear to form neural progenitors and migratory NC cells, while adopting a more posterior identity over time.

### NMP-derived cells stabilise as epithelial pre-neural progenitors

To determine the extent to which NMP-derived cells undergo differentiation, epithelial and mesenchymal cells were enzymatically separated at P5, profiled by bulk RNA-Seq, and compared with the original NMP-like transcriptional profiles (Figure S6A). The temporal progression from 36h to P5 accounted for the majority of gene variation (PC1, ~70%) that was detected. The lineage bifurcation of NMP descendants led to the identification of 907 differentially expressed genes between epithelial and mesenchymal cells (426 genes up in epithelial and 481 genes up in mesenchymal cells; FDR <1%, ≥2-fold change, DESeq2 base mean >100 reads—supplementary file 4). Strikingly, the enrichment analysis of upregulated genes for cellular component GO terms showed that epithelial and mesenchymal cells were linked to key attributes of nerve cell differentiation (e.g. ‘synapse’ and ‘axon’) and NC cell migration (e.g. ‘extracellular matrix’ and ‘adherens junction’), respectively (Figure 4A,B, supplementary file 4). However, we did not observe expression of post-mitotic neuronal markers such as ELAV-like RNA Binding protein 3/4 (ELAVL3/4) or Tubulin-beta class III (TUBB3) suggesting cells are of an immature neuronal cell type with no synapses and axons yet (Delile et al., 2019). Molecular function GO terms for both samples were similar, and primarily reflected the large number of transcription factors expressed, but also included ‘growth factor binding’ terms which represented WNT/FGF signalling genes in addition to TGF-β superfamily signalling genes (Supplementary file 5). Few of these genes were differentially expressed between epithelial and mesenchymal samples, and they included both positive (BMP4/5/7) and negative (GREM1 and CER1) regulators of TGF-β signalling (Figure S6B, Supplementary file 5). Together this analysis further suggests that the epithelial cells, unlike the mesenchymal cells, are a neuronal cell type and that endogenous signalling pathways, including the TGF-β superfamily, may influence cell identity over time.

**Figure 4:**
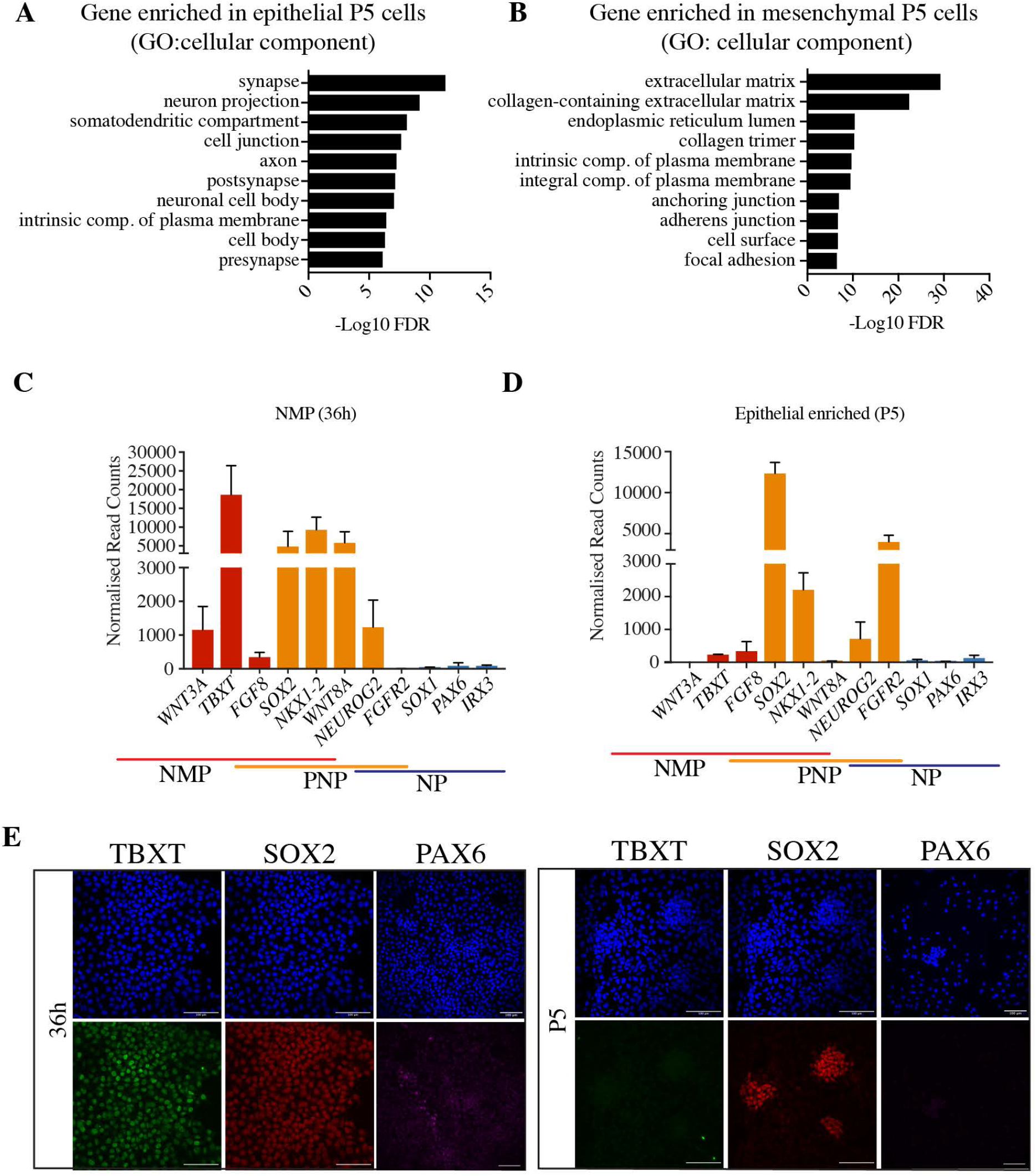
NMP-derived cells stabilise as epithelial pre-neural progenitors. A,B) Graphs showing cellular component GO analysis for differentially expressed genes in P5 epithelial samples (A) and P5 mesenchymal samples (B). The corresponding Benjamini and Hochberg adjusted p-values (FDR) are shown. C, D) Normalised expression levels of known markers of NMPs (*WNT3A*, *TBXT*, *FGF8*, *SOX2*, *NKX1-2* and *WNT8A/C*), PNPs (*SOX2*, *NKX1-2*, *WNT8A/C, NEUROG2 and FGFR2*) and NPs (*PAX6*, IRX3, FGFR2, *NEUROG2* and *SOX1*) at 36h (C) and in P5 epithelial colonies (D) as determined by RNA-seq. Error bars show SEM (n = 3 biological replicates). E) Representative immunostaining of TBXT (green), SOX2 (red) and PAX6 (magenta) confirming the expression patterns shown in (A and B). Scale bars, 100 μm.

Next, a panel of previously established NMP, PNP and neural progenitor marker genes were used to pinpoint neural progression *in vitro* (Verrier et al., 2018, Ribes et al., 2008, Olivera-Martinez et al., 2014). As expected, 36h cells were positive for NMP markers (*FGF8*, *WNT3A* and *TBXT*) and NMP/PNP (*SOX2*, *NKX1-2* and *WNT8A/C*), while the NP determinants *PAX6, IRX3* and *SOX1* were hardly transcribed (Figure 4C). By P5, epithelial cells had lost most NMP-exclusive expression, while the PNP markers *SOX2* and *NKX1-2* were retained (Figure 4D). *NEUROG2* and FGFR2, two PNT/NT markers, were also active in P5 epithelial cells (Ribes et al., 2008, Olivera-Martinez et al., 2014). Furthermore, neural progenitor markers were low or absent in epithelial P5 cells (Figure 4D). Immunofluorescence for TBXT, SOX2 and PAX6 confirmed this transcriptional analysis, some of which was further validated by RT-qPCR (Figure S6C). Together, we find that epithelial colonies have a PNP identity and do not express key neural maturation genes.

### NMP-derived mesenchymal cells are NC

We next sought to determine the identity of the mesenchymal cells. *In vitro* studies have revealed that NMPs can become trunk NC cells, a migratory mesenchymal cell population which goes on to form tissues including cartilage, bone and smooth muscle (Frith et al., 2018, Hackland et al., 2019, Leung et al., 2016). Moreover, our bulk RNA-Seq suggested that over passaging there was an increase in genes associated with cell migration and NC derivatives, concomitant with the reduction of epithelial cells and increase of differentiating mesenchymal cells in late passages (Figures 2E and 3A,B). Thus, we first determined whether mesenchymal P5 cells had acquired NC-specific gene expression. Transcriptome-wide analysis showed that several NC markers genes, including *SNAI1, SOX9* and *SOX10,* were significantly higher in mesenchymal cells compared with their epithelial PNP counterparts (Figure 5A,B). This was corroborated by immunofluorescence of P5 tissue cultures, which showed SNAI1^+^ and SOX10^+^ mesenchymal cells scattered between SOX2^+^/CDX2^+^ PNP colonies (Figures 2B and 5C). In support of a posterior NC identity, mesenchymal P5 and P8 cells progressively expressed more posterior HOX genes, mirroring the PNP rostrocaudal identity (Figure 3C). By contrast, the cranial NC marker ETS1 was only detectable in a few mesenchymal cells (Figure 5D). To determine if mesenchymal cells were capable of generating trunk NC derivatives, mesenchymal P5 cells were exposed to 1% fetal calf serum (FCS) for 7 days to convert them into NC-derived vasculature smooth muscle, containing cytoplasmic fibres of α-smooth muscle actin (α-SMA also known as ACTA2; Figure 5E,F) (Mohlin et al., 2019). Together, these results show that the mesenchymal cells surrounding PNPs are functional posterior NC cells.

**Figure 5:**
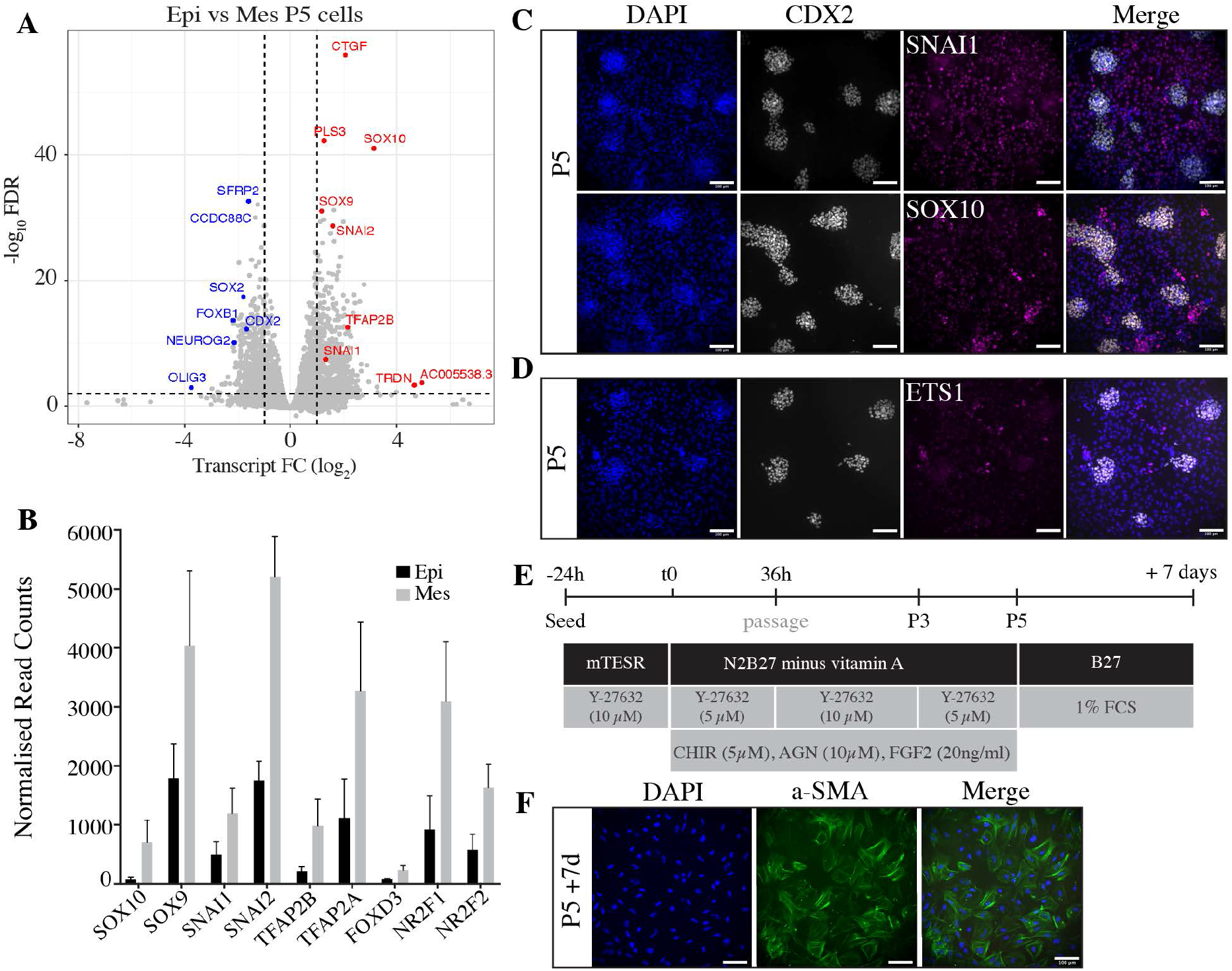
Mesenchymal cells have a neural crest identity. A) Volcano plot showing differential expression between epithelial and mesenchymal cell at P5. Significant genes are highlighted in blue (epithelial) and red (mesenchymal). B) Normalised expression levels of known markers of NC genes (*SOX10, SOX9, SNAI1, SNAI2, TFAP2B, TFAP2A, FOXD3, NR2F1* and *NR2F2*) which are significantly upregulated in mesenchymal enriched samples compared to epithelial as determined by RNA-seq. Error bars show SEM (n = 3 biological replicates). (C,D) Representative immunostaining of NC markers SNAI1, SOX10 (C) and ETS1 (D), co-stained with epithelial PNP marker CDX2 (grey) and the nuclear stain DAPI (blue). Scale bar, 100 μm. E) Scheme for generating NMP/PNP-derived NC derivative smooth muscle. F) Representative immunostaining of α-SMA (green) and nuclear stain DAPI (blue) in NMP/PNP-derived vasculature smooth muscle cells. Scale bar, 100 μm.

### NMP-derived trunk PNPs are stem cell-like and give rise to migratory NC

The immunofluorescence analysis of fixed PNP/NC cell cultures revealed that some nuclei found within tightly clustered PNP colonies were negative for CDX2, but positive for SNAI1, suggesting that they are undergoing epithelial-to-mesenchymal transition (EMT) and becoming NC cells (Cano et al., 2000, Simoes-Costa and Bronner, 2015) (Figure 6A, 2B,E). To test this idea, PNP colonies (CDX2^+^/SNAI1^−^) purified from NC cells using selective detachment were sub-cultured for four passages (P+1 to P+4) (Figure 6B). Immunofluorescence staining showed that, despite the low percentage of SNAI1^+^ NC (8%) cells in P+1 cultures, by P+4 40% of the cells were CDX2^−^/SNAI1^+^ suggesting that PNPs undergo EMT to generate NC cells (Figure 6B,C). To exclude the possibility that after PNP purification, the remaining NC cells repopulate the culture over passaging, single cells from the PNP or NC enriched samples were re-plated by fluorescence-activated cell sorting (FACS) into single wells (Figure S7A). No colonies arose from single NC cells, suggesting that these cells have limited proliferative capacity. By contrast, single PNPs gave rise to clonal cell lines which consisted of epithelial colonies (CDX2^+^/SOX2^+^), and surrounding NC cells (Figure 6D,E). Thus, the PNPs showed stem cell-like behaviour by undergoing self-renewal and differentiating into NC cells.

**Figure 6:**
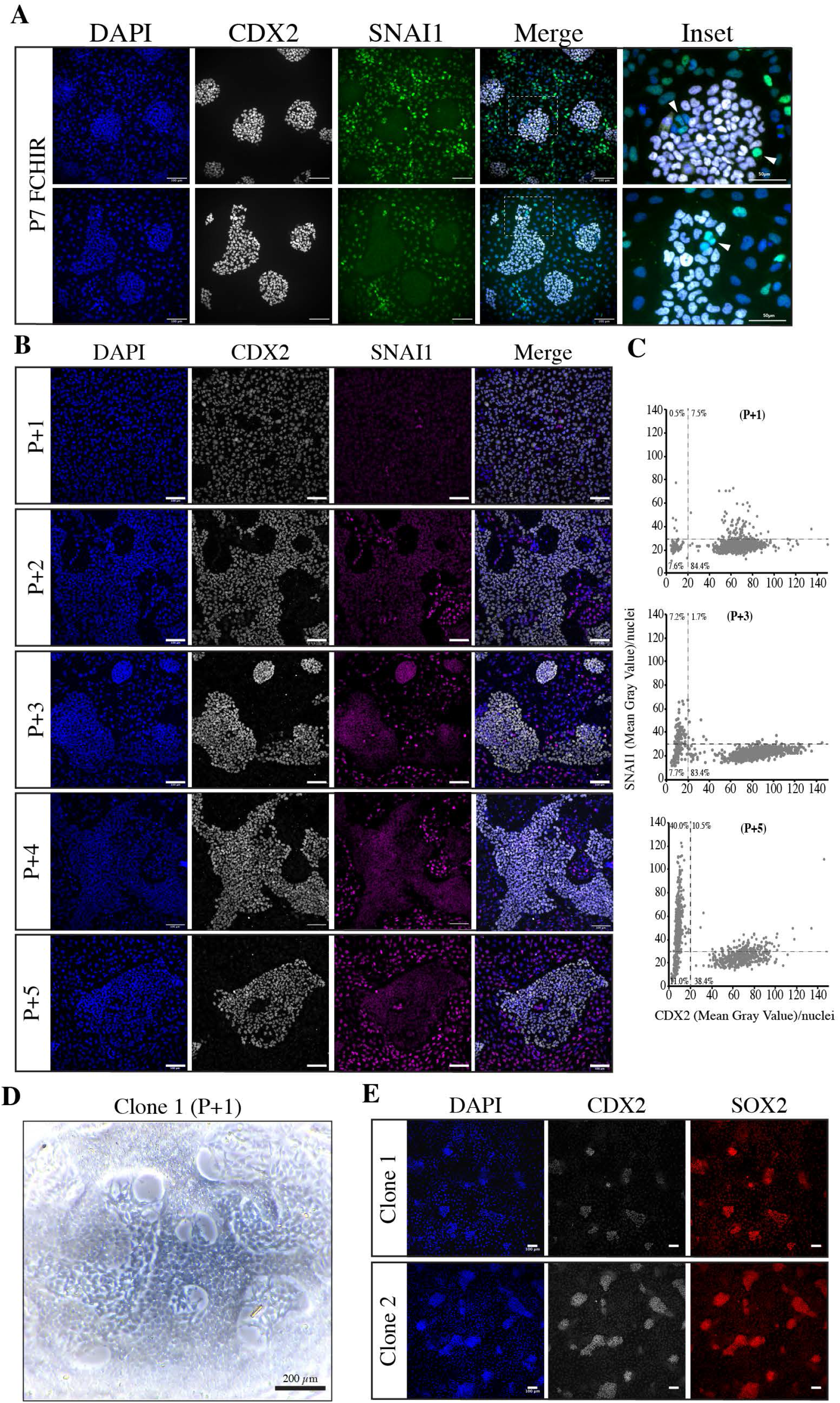
Epithelial PNPs give rise to migratory neural crest cells. A) Representative immunostaining of CDX2 (grey) and SNAI1 (green) co-stained with nuclear stain DAPI (blue) in P7 PNP/NC cultures. Inset shows magnified region identified by white dashed line and arrow marks examples of CDX2^−^/SOX2^−^/SNAI1^+^nuclei within PNP clusters. Scale bars, 100μm or 50 μm (inset). B) Representative immunostaining of CDX2 (grey), SNAI1 (magenta) and nuclear stain DAPI (blue) in epithelial P5 cells which were serially passaged for four passages (P+1 to p+4) following selective detachment enrichment. (C) Dot plot showing the mean grey value/nuclei of CDX2 and SNAI1 at P+1, P+3 and P+4 panels shown in (B). Each graph shows >900 nuclei. D) Representative bright-field image of a sub-clone generated from the epithelial enriched fragment after 1 passage. Scale bar, 200 μm E) Representative immunostaining analysis of CDX2 (grey), SOX2 (red) and nuclear stain DAPI (blue) in two independent sub-clones generated from the epithelial enriched samples after serial 4 passages. Scale bar, 100 μm.

### Modulation of TGF-β and SHH signalling locks in PNP rostrocaudal axis information by preventing trunk-to-tail transition

We have shown that the combined modulation of Wnt/FGF and RA signalling generated posterior PNPs. However, transcriptomics and lineage analysis indicated that PNP maintenance may be compromised by NC bifurcations, the progressive activation of more posterior HOX genes, and late-passage differentiation/cell death. In line with this, a known regulator of trunk-to-tail transition and terminal HOX induction, GDF11 was found to be significantly up-regulated in late passages compared to early passages (Figure 7A). Increased *GDF11* expression preceeds activation of the terminal *HOX13* genes and coincides with the down-regulation of the stem cell marker *LIN28A*, leading to a loss in cell proliferation (Aires et al., 2019, Jurberg et al., 2013, Robinton et al., 2019)(Figure 7B,C). As such, in an attempt to prevent this progressive posteriorisation and NC commitment, we supplemented our culture medium with modulators of the TGF-β pathway (Figure 7D). Inhibitors of Activin/Nodal (SB431542, SB) and BMP (LDN193189, LDN) signalling were used to supress TGF-β and NC specification (Inman et al., 2002, Cuny et al., 2008, Halder et al., 2005, Das et al., 2009, Liem et al., 1997, Stuhlmiller and Garcia-Castro, 2012). Furthermore, to mimic signals that arise from the notochord during neural tube folding/cavitation and induce a ventral identity in differentiated neuronal cultures, a smoothened agonist (SAG) was used to stimulate Sonic Hedgehog (SHH) signalling (Sasai et al., 2014, Jessell, 2000).

**Figure 7:**
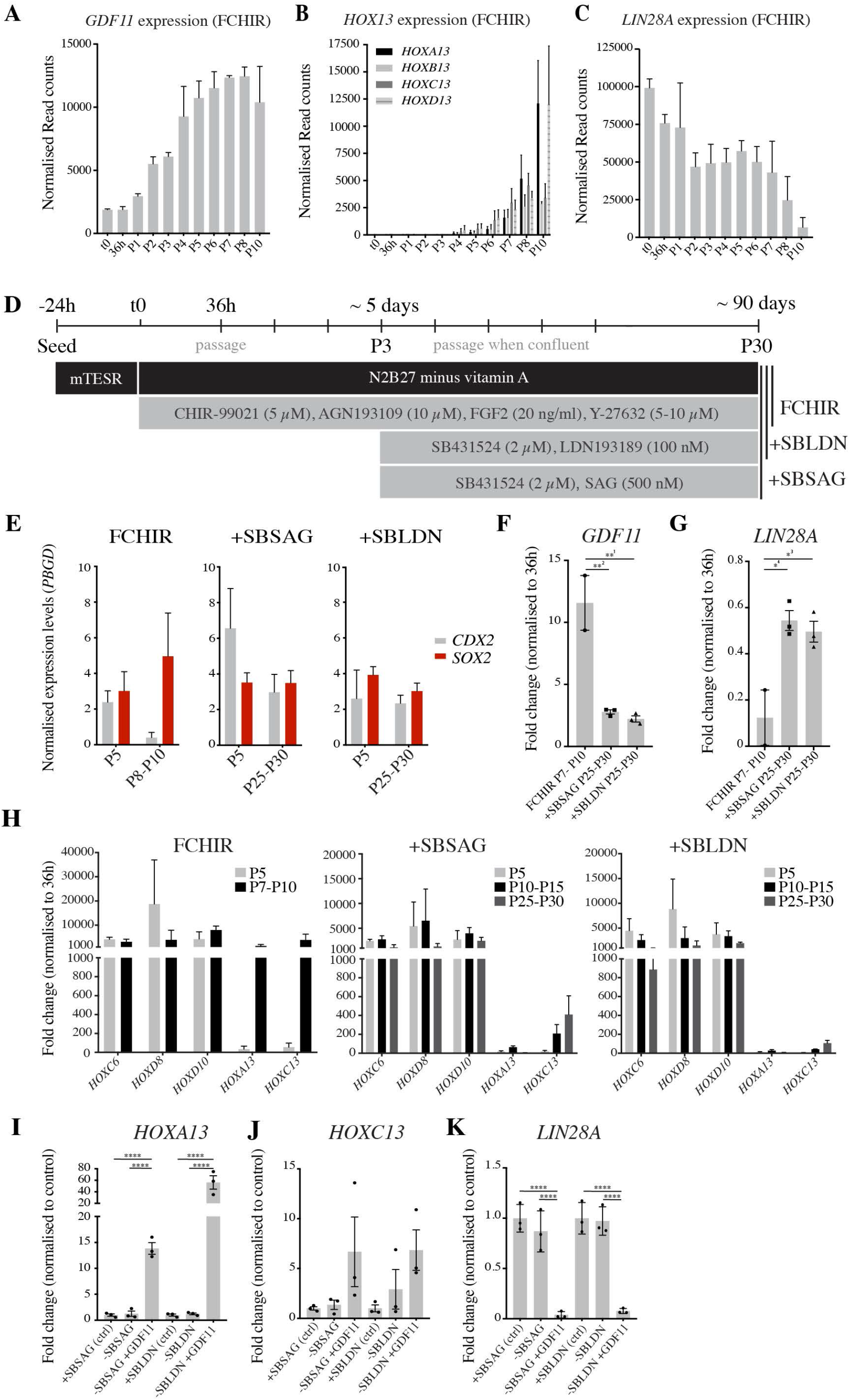
Modulation of TGF-β and SHH signalling locks in A/P information. A,B,C) Normalised expression levels of *GDF11* (A), HOX13 (B) and LIN28A (C) at each passage as determined by RNA-seq. Error bars show SEM (n = 3 biological replicates). D) Scheme for generating and maintaining PNPs. At passage 3 either SB and LDN (+SBLDN) were added, or SB and SAG (+SBSAG) were added to the standard medium (FCHIR). E) Transcriptional quantification (RT-qPCR) of CDX2 and SOX2 at early (P5) and later passages (FCHIR; P8-P10 and +SBLDN and +SBSAG; P25 −P30). Expression levels normalised to the reference gene *PBGD.* Error bars show SEM (n = 2-5). F, G) Transcriptional quantification (RT-qPCR) of *GDF11* (F) and *LIN28A* (G) shown by fold change over 36h and normalised to the reference gene *PBGD* in late passages PNPs (FCHIR; P8-P10 and +SBLDN and +SBSAG; P25-P30). Error bars show mean with SEM (n = 2/3). **P^1^ = 0.0019, **P^2^ = 0.0014, *P^3^ = 0.0107, *P^4^ = 0.0174 (ANOVA) H) Graphs showing the transcriptional quantification (RT-qPCR) of selected *HOX* genes at early (P5) and mid (FCHIR; P8-P10 and +SBLDN and +SBSAG; P10-P15) AND late (+SBLDN and +SBSAG; P25-P30) in all conditions tested as indicated in (D). Expression levels are presented as fold change over the 36h time point and were normalised to the reference gene *PBGD.* Error bars show mean with SEM (n = 2/3). I,J,K) Transcriptional quantification (RT-qPCR) of *HOXA13* (I) and *HOXC13* (J) and *LIN28A* (K) in +SBLDN or +SBSSAG (ctrl) conditions, +SBLDN or +SBSAG without SB, LDN or SAG (-SBLDN/-SBSAG) and +SBLDN or +SBSAG without SB, LDN or SAG but with GDF11 (-SBLDN/-SBSAG +GDF11). All experiments were completed with P25-P30 cultures. Expression levels normalised to the reference gene *PBGD.* Error bars show SEM (n = 2-5). ****P<0.001 (ANOVA).

The combined addition of SB and LDN (+SBLDN) or SB and SAG (+SBSAG) at P3 resulted in stabilisation of PNPs for over 30 passages (90 days). At early passages (P5/P6), the addition of small molecules did not compromise the formation of CDX2^+^/SOX2^+^ PNPs, which organised into typical tightly associated colonies surrounded by loosely packed SNAI1^+^ cells. (Figure S8A,B,C). However, both supplemented conditions modestly increased the percentage of SOX2^+^/CDX2^+^ cells as quantified by flow cytometry in late passages (P9/P10) (Figure S8C,D). Cells maintained in +SBSAG and +SBLDN had significantly prolonged *CDX2* and *SOX2* gene expression for up to 30 passages (Figure 7E). Based on the transcriptional profiling of *HOX* genes, the positional value of the PNPs was locked at the thoracic level, considerably slowing down the upregulation of terminal *HOXC13* and *HOXA1*3 (Figure 7H). As expected, in comparison to P7-P10 FCHIR generated cells, *GDF11* expression was significantly lower in +SBSAG and +SBLDN cultures (Figure 7F). In line with this, *LIN28A*, which is known to be down-regulated in response to *HOX13* expression, was considerably reduced in FCHIR cultures by P7-P10 (Aires et al., 2019) (Figure 7G). To test if the trunk-to-tail transition can be induced in PNPs after long-term TGF-β inhibition, we added exogenous human recombinant GDF11 to P28-P30 cultures for 48-72h. This short term treatment of GDF11 was sufficient to induce *HOXA13* and *HOXC13* gene expression supress *LIN28A* expression (Figure 7I,J,K).

Notably, Verrier et al. (2018) also used dual inhibition of Nodal/Activin and BMP signals to generate RA-induced neural progenitors from NMPs. However, these cells were not maintained over long time periods presumably because of their exposure to RA. In our tissue cultures, the RA target *PAX6* remained silent in +SBLDN or +SBSAG addition at P6/7 (Figure S8E). These results therefore, show that PNPs can be locked in a thoracic identity and grown in culture for long periods of time via the addition of TGF-β inhibitors by preventing the GDF11/LIN28A-mediated trunk-to-tail transition.

### PNPs can give rise to spinal cord neurons

To establish the neuronal potential of RA-deprived PNPs, we terminally differentiated P5 FCHIR and P25 +SBSAG/+SBLDN long-term PNPs into neurons (Figure 8A). Analysis of lateral motor column (LMC; FOXP1), dorsal interneuron/lateral motor column marker (LHX1) and medial motor column markers (MMC; LHX3) found that all PNP conditions preferentially generated LHX1^+^/ TUJ^+^ cells although did not express ISL1 (Figure 8B,D). The prescense of LHX1^+^/ISL1^−^positive neurons, suggests neurons may be lateral LMC (LHX1^+^/ISL2^+^), interneurons of the p2-dp2 domains or medial LMC which no longer express early motor neuron markers (Francius and Clotman, 2014, Zannino and Sagerström, 2015) (Figure 8B). Few cells were found to express LHX3 indicating cells preferentially differntate MMC motor neurons (Figure 8C). Furthermore, more CHX10^+^ cells were noted in +SBSAG PNP-derived cultures, suggesting SHH signalling may introduce a more ventral identity after differentiation giving rise to V2a interneurons (CHX10^+^/TUJ^+^) although sustained SHH signalling throughout early differentation should increase the yield of ventralised neurons (Figure S9A) (Thaler et al., 2002, Clovis et al., 2016, Le Dréau and Martí, 2012). Together, these results show that our PNPs can generate various spinal cord derivatives demonstrating neuronal potential.

**Figure 8:**
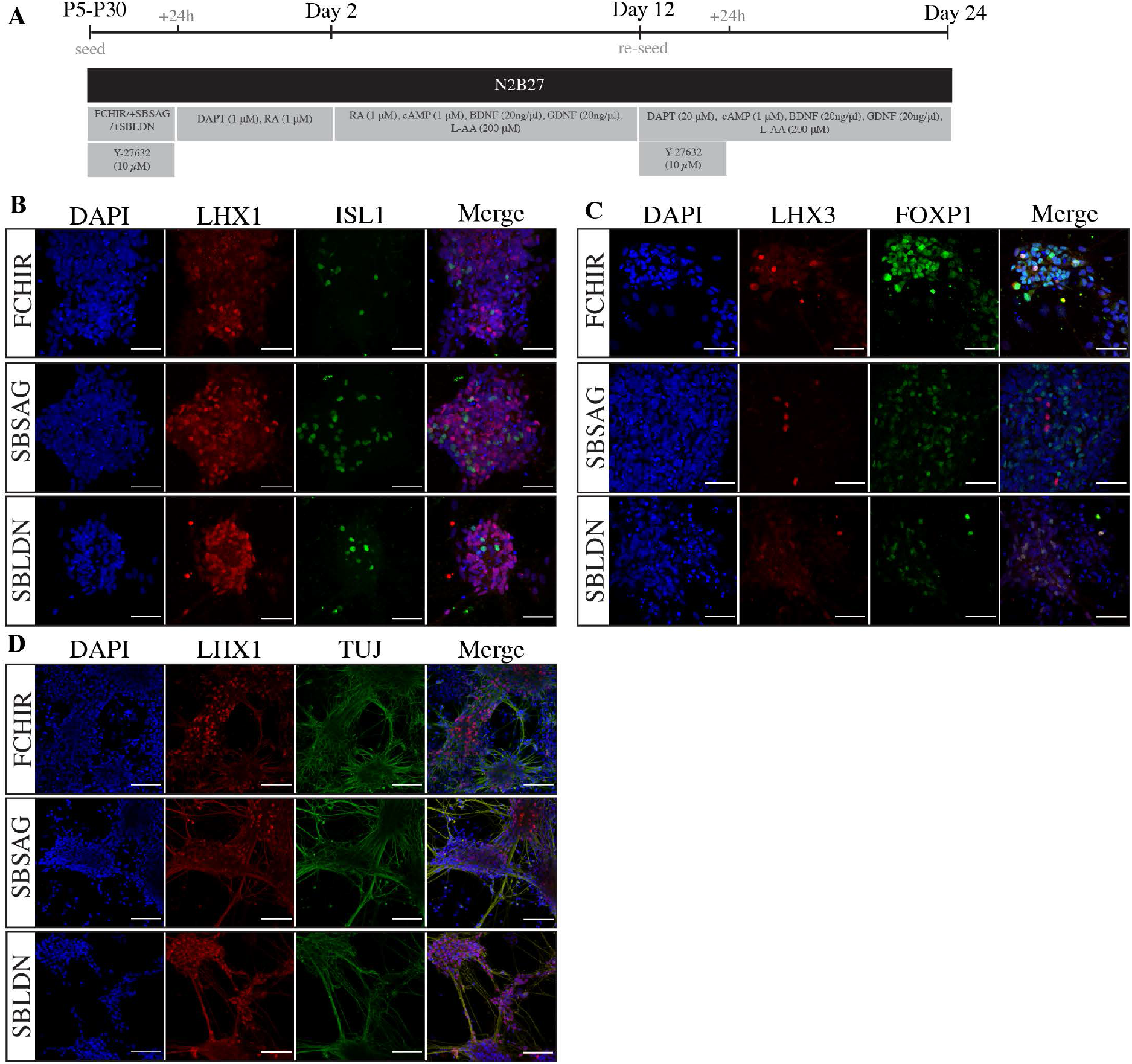
PNPs can be differentiated into neural derivatives. A) Scheme for generating differentiated neuronal cultures. PNPs are dissociated and plated at low density and then exposed to neural inducing factors shown. B,C,D) Representative immunostaining of differentiated neuronal cultures showing (B) LIM homeobox 1 (LHX1, red) and Islet1 (ISL1, green) or (C) LIM homeobox 3 (LHX3, red) and FOXP1 (red). Nuclei were stained with DAPI (blue) and (D) LHX1 (red) paired with βIII-tubulin (TUJ, green) Scale bars, 100μm.

## DISCUSSION

The NMP niche is maintained by Wnt/FGF-mediated autoregulatory loops, CYP26A1-mediated RA signal suppression, and active RARγ–mediated transcriptional repression (Janesick et al., 2014, Koide et al., 2001, Sakai et al., 2001, Yamaguchi et al., 1999, Deng et al., 1994, Cunningham et al., 2015, Takemoto et al., 2006, Martin and Kimelman, 2008, Takada et al., 1994, Liu et al., 1999, Abu-Abed et al., 2001, Martin and Kimelman, 2010). By simultaneously controlling these multiple signalling pathways *in vitro*, we have generated regionalised spinal cord progenitors and NC cells from hPSCs. Our protocol consistently yields a well-defined population of spinal cord PNPs and NC cells different rostrocaudal identities, providing a valuable source of spinal cord cells and NC which hold the potential for drug screening, detailed disease modelling, or therapeutic applications. Moreover, our model provides a robust platform to study cellular commitments and transitions within the developing human spinal cord at greater detail. In the long term, we hope that use of our protocol will improve the understanding of selective neuronal vulnerability, a recognised, yet poorly understood feature of neurodegenerative disease and spinal cord injury. More recently, Wind et al., (2020) similarly showed that prolonged FGF/Wnt signalling can generate spinal cord neural progenitor populations capable integrate with into the chick neural tube, suggesting their potential of the cells for disease modelling and regenerative therapy.

Since Gouti et al., (2014) showed that trunk neuronal derivatives can be generated from mouse NMP-derived cells, various protocols have emerged for generating spinal cord progenitors from hPSCs. However, these often do not produce progenitors which undergo complete rostrocaudal diversification and therefore do fully mimic *in vivo* development. Furthermore, previous studies showed that *in vitro* generated NMP-like cells, when passaged back into FGF/ CHIR or CHIR alone, commit to a mesodermal lineage (Gouti et al., 2014, Turner et al., 2014). In contrast, we observed a gradual decrease in TBXT and TBX6 expression, and commitment of NMP-like cells to a neural trajectory. Interestingly, this commitment appeared to occur independently of RA signalling. More recently, Edri *et al.*, (2019) found NMP-like cells derived rather than ESCs, resemble more accurately their counterparts *in vivo* and have a similar progressive commitment to a neural fate after passaging (Edri et al., 2019). This is in line with our observations using hPSCs, and could reflect that hPSCs correspond more closely to the primed pluripotency state of mouse EpiSCs (Nichols and Smith, 2011, Brons et al., 2007). Compared to previous studies of *in vitro*-derived NMPs, the PNPs reported here are a more stable source of spinal cord as their maintenance does not depend on the delicate balance between TBXT and SOX2, and they do not express critical RA-responsive neurogenic genes like PAX6 and SOX1, which would promote their fate progression to spinal cord neurons (Gentsch et al., 2017, Janesick et al., 2015).

While PNPs were efficiently derived from NMPs, their long-term maintenance was accompanied by progressive posteriorisation and NC delamination. Thus, to promote PNP self-renewal, we tried to mimic the niche environment of axial stem cells by inhibiting TGF-β and stimulating SHH signalling. Furthermore, TGF-β superfamily signalling members GDF11/GDF8 are known to promote trunk-to-tail transition, resulting in the up-regulation of HOX13 genes and the loss of LIN28A, a key factor, for the proliferation of tail bud progenitors (Aires et al., 2019). TGF-β signal inhibition favoured PNP fate over time and locked PNPs in a thoracic HOX identity for up to 30 passages, highlighting the importance of TGF-β signal inhibition in maintaining trunk PNPs. Previous *in vivo* data supports this observation, as the inhibitory TGF-β signal transducer *SMAD6*, is specifically expressed in the PNT, while the Activin-neutraliser Follistatin (FST) is required for dorsal-ventral patterning and neuronal fate specification in response to SHH signalling (Olivera-Martinez et al., 2014, Liem et al., 2000). Specifically, our data indicates that ALK4, ALK5 and ALK7 inhibition by SB431542 is acting to prevent GDF11 signalling and is sufficient to promote PNP identity and viability in our culture by maintaining LIN28A expression (Andersson et al., 2006). This is in line with recent *in vivo* evidence describing the involvement of TGFBR1/GDF11 in secondary neurulation (Dias et al., 2020, Aires et al., 2019).

Our work also established that PNPs undergo EMT to form NC cells with corresponding rostrocaudal identity. Recent studies have shown that cranial NC is specified at the neural plate border and trunk NC arises from the NMP niche in a BMP dependent manner (Frith et al., 2018, Wymeersch et al., 2016, Stuhlmiller and Garcia-Castro, 2012). Surprisingly, the addition of the BMP inhibitor (LDN) did not prevent NC specification in long-term PNPs. BMP-mediated inhibition of NC was possibly GDF11 inhibition, which in mouse was found to lead to an increase in SOX10, suggesting an increased specification of NC specification in the tail (Aires et al., 2019). As a result, it is unclear why BMP inhibition does not prevent NC specification but we hypothesise that incomplete BMP signalling inhibition is the most likely candidate driving NC specification in this setting, with only an intermediate level of BMP signalling required to induce NC commitment (Frith et al., 2018, Hackland et al., 2017). Further work to test this hypothesised is required. Furthermore, we did not observe any direct NC specification from NMPs indicating NC specifcation *in vivo* may occur at the PNT. However, it remains a possibility that NC can be specified from NMPs and PNPs, and will be interesting to explore further.

In conclusion, we show with sustained Wnt/FGF signalling PNPs undergo collinear HOX gene expression and the transition to a pre-neural fate (Mouilleau et al., 2020, Edri et al., 2019, Wind et al., 2020). RA inhibition prevents the uprgulation of RA-responsive neural determinants genes such as PAX6 and maintains expression of genes associated with a PNP idenity. We further suggest, based on previous studies and the generation of spinal cord neurons, that removal of FGF/Wnt signalling and addition of RA signalling permits differentiation to neural progenitors and an upregulation of neurogenic genes (Figure 9) (Verrier et al., 2018, Diez del Corral et al., 2002, Diez del Corral et al., 2003, Wind et al., 2020). Furthermore, single PNPs undergo ‘self-renewal’ due to high *LIN28A* and low *HOX13* expression until PNPs undergo trunk-to-tail transition as a result of increased GDF11 signalling. The addition of TGF-β inhibition combined with BMP inhibition or SHH agonism (+SBLDN/+SBSAG) prevents GDF11 upregulation and subsequent loss of LIN28A, resulting in stabilisation of PNPs in a thoracic identity for up to 30 passages. Finally, PNPs give rise to NC as they progress through a rostro to caudal identity, the first protocol to our knowledge to generate NC *in vitro* with diverse rostrocaudal identity.

**Figure 9:**
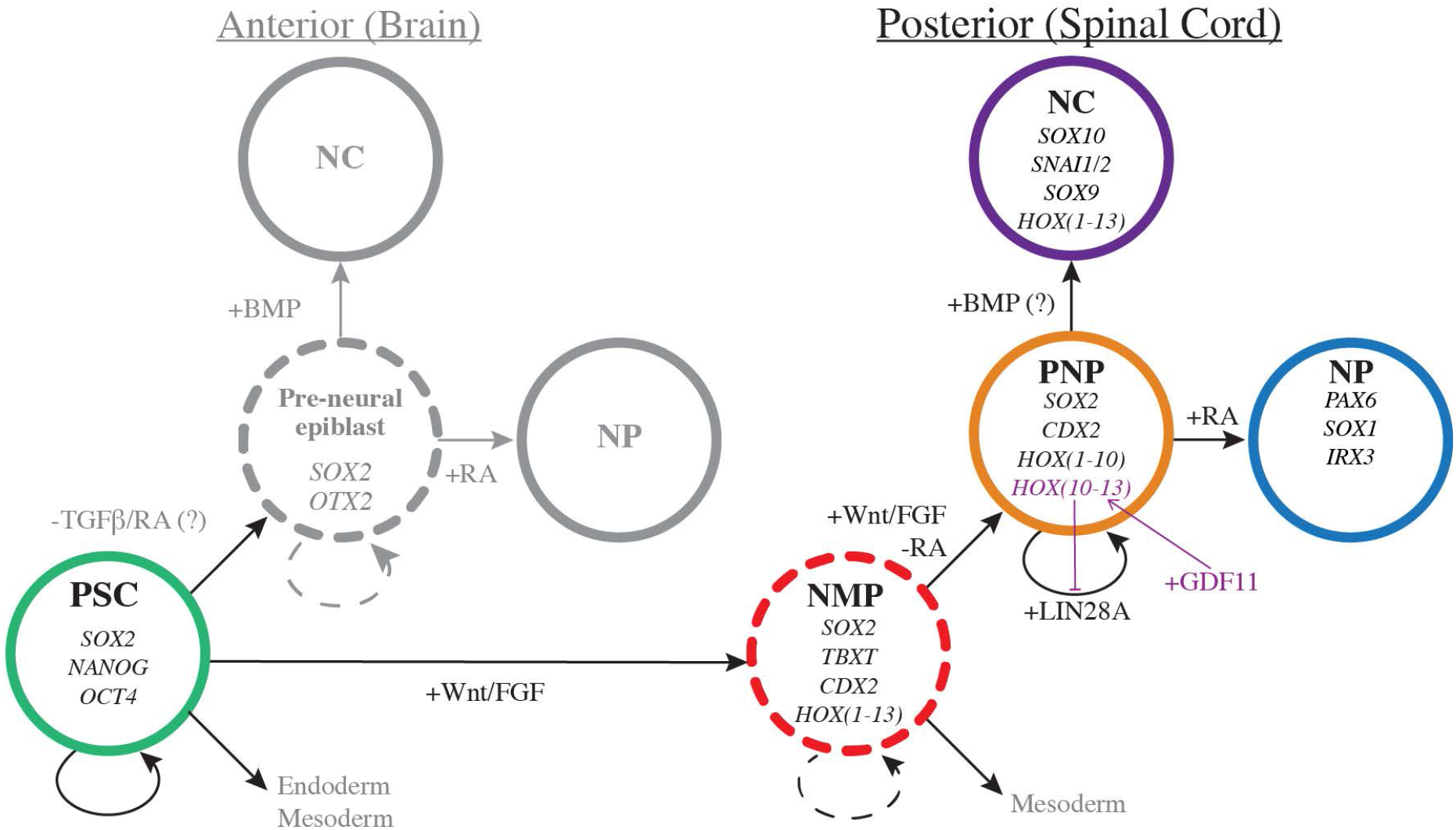
NMP-derived PNPs self-renew, give rise to trunk NC or can be differentiated to neurons. Diagrammatic model summarising the generation of anterior (brain) and posterior (spinal cord) neural progenitors *in vitro*. When treated with inhibitors of TGF-β signalling pluripotent stem cells (PSC) in the give rise to a transient pre-neural epiblast state, which in turn give rise to anterior NC and neural progenitors (NP) of the brain. NMPs, which give rise to posterior neural tissue, are generated from PSC in response to Wnt/FGF signalling. In sustained Wnt/FGF signalling and in the absence of RA, NMPs to differentiate to a PNP intermediate which are able to self-renew and give rise to neural progenitors or NC in the presence of RA or BMP, respectively. GDF11 signaling results in upregulation of terminal HOX genes and results in loss of LIN28A ultimately leading to loss of PNP self renewal. Transient cell states are shown using dotted lines and cells with self-renewal capacity are shown with curved arrows.

## ACKNOWLEDGEMENTS

We thank members of the following scientific platforms and units of the Francis Crick Institute for their expertise, support and use of the facilities: advanced sequencing facility, advanced light microscopy facility, the human embryo and stem cell unit, bioinformatics and biostatistics and research illustration and graphics. We also thank Rickie Patani, Jamie Mitchell, James Briscoe, Vicki Metzi, Teresa Rayon, Alessia Caramello, Robin Lovell-Badge, Siew-Lan Ang and Francois Guillemot for advice, help and reagents; Rebecca Jones and Clara Collart for critical reading of the manuscript; and the Smith lab for discussions and advice.

## CONTRIBUTIONS

FC: Conceptualization, Validation, Methodology, Investigation, Formal analysis, Writing—original draft preparation, Supervision, Project administration

GEG: Conceptualization, Methodology, Investigation, Supervision, Project administration, Writing— review & editing

RM: Software, Methodology, Formal analysis, Writing—review & editing CB: Investigation, Writing—review & editing

LH: Methodology, Investigation, Resources, Writing—review & editing AHR: Investigation

JCS: Conceptualization, Writing—review & editing, Supervision, Funding acquisition

ASB: Conceptualization, Methodology, Investigation, Writing—review & editing, Supervision, Project administration, Funding acquisition

## METHODS

### Human pluripotent stem cell culture

Human ESCs (WA09 and WA01, WiCell) and human iPSCs (AICS-23, Allen Institute) were maintained in feeder-free cultures, plated on Corning Matrigel Growth Factor Reduced (GFR) Basement Membrane Matrix (Corning Incorporated, 354230) and grown in mTESR1 (STEMCELL technologies, 85850). Cells were passaged as aggregates at a ratio of 1:10/15 using Gibco Versene Solution (Thermo Fisher Scientific, 15040066). All experiments were completed within 15 passages after recovery from cryopreservation and screened for mycoplasma monthly. Prior to cryopreservation, hPSCs were assessed for genetic stability by KaryoStat and indicators of pluripotency were assessed by PluriTest (Thermo Fisher Scientific). hPSCs were subject to routine pluripotency using BD Stemflow Human and Mouse Pluripotent Stem Cell Analysis Kit (BD Biosciences, 560477) as recommended by the manufacturers, or by immunostaining against OCT3/4, SOX2 and NANOG (see Table S5 for antibody details) using the standard immunostaining protocol below. All experiments with hESCs were approved by the UK Stem Cell Bank steering committee (SCSC13-03).

### NMP differentiation

For differentiation into NMPs, confluent hPSCs were dissociated into single cells using Gibco TrypLE Express (Thermo Fisher Scientific, 12604013) and plated at a density of 50,000 cells/cm^2^ on Matrigel hESC-Qualified Matrix (Corning Incorporated, 354277). Cells were plated in mTESR1 supplemented with 10 μM Y-27632 (Tocris, 1254) for a 24h to 36h to allow recovery before starting differentiation into NMPs. Following recovery time, cells were grown in Dulbecco’s Modified Eagle Medium/Nutrient Mixture F-12 (DMEM/F-12, Thermo Fisher Scientific, 10565018) supplemented with 1x Gibco B-27 supplement minus vitamin A (Thermo Fisher Scientific, 12587010) and 1x Gibco N2 (Thermo Fisher Scientific, 17502048), 4-6 μM CHIR-99021 (Selleck Chem, S2924-SEL-5mg), 10 μM AGN193109 sodium salt (Santa Cruz, sc-210768) and 20 ng/ml FGF2 (R&D systems, 233-FB-025) referred to from now on as NMP differentiation medium. NMP differentiation medium was supplemented with and 5 μM Y-27632 (Tocris).

### PNP long term culture

To generate PNPs, NMPs were passaged at 36h using TrypLE express (Thermo Fisher Scientific) and when confluent thereafter. Cells were passaged as single cells at a ratio of 1:6 into NMP differentiation medium, supplemented with 10 μM Y-27632 (Tocris). During passage 1 to 3 progenitors were found to detach from the dish forming spheres. If this occurred, spheres were dissociated into single cells and re-plated immediately. PNP generation was more successful if cells did not detach, therefore, to prevent cells detaching during this period cells were passaged before reaching high confluency. In addition, cells were only removed from the 37oC incubator when ready to passage, as the temperature fluctuations promoted detachment. From passage 3 cells were grown NMP differentiation medium supplemented with 5 μM Y-27632 (Tocris). Human iPSCs were found to detach more readily than hESCs. PNPs could be maintained, for 8 to 12 passages using standard conditions as above, passaging every 3-4 days when 80-90% confluent. To lock A-P axis progression, 2 μM SB431542 (CELL guidance systems, SM33-10) and 100 nM LDN193189 (Sigma-Aldrich, SML0559-5MG) or SB431542 (CELL guidance systems, SM33-10) and 500 nM smoothened agonist (SAG, Sigma-Aldrich, 566660-1mg) were added to NMP differentiation medium at passage 3. For selective detachment, 90% confluent PNPs were washed with PBS and treated with TrypLE express (Thermo Fisher Scientific) at 37oC for 3-5 mins. When mesenchymal cells started to detach, cells were gently removed by tilting the plate side-to-side. TrypLE containing the detached mesenchymal cells was carefully removed. Remaining epithelial cells were washed off the vessel using basal medium.

### Neuronal differentiation

To generate neurons, we used a modified protocol based on a previously published neural differentiation protocol (Lippmann et al., 2015). 80-90% confluent PNP/NC cultures were dissociated to single cells and plated at 33,000 cells/cm^2^ onto Matrigel hESC-Qualified matrix (Corning) into the applicable former culture medium (NMP differentiation medium plus or minus SBLDN or SBSAG). 24h after plating, medium was replaced with neural differentiation medium consisting of Gibco neural basal medium (Thermo Fisher Scientific, 21103049) supplemented with Gibco 1x B27 supplement (Thermo Fisher Scientific, 17504044) and 1x N2 (Thermo Fisher Scientific), 2 μM DAPT (Chem Cruz, sc-201315) and 1 μM retinoic acid (RA, Sigma Aldrich, sc-210768) for 48h. Following 48h treatment, media was replaced with 10 ng/ml brain-derived neurotrophic factor (BDNF, PeproTech, 450-02-2UG), 10 ng/ml glial-derived neurotrophic factor (GDNF, PeproTech, 450-10-2UG), 1 μM retinoic acid (RA, Sigma Aldrich, sc-210768), 1 μM cAMP (Sigma Aldrich, A6885-100mg) and 200 μM L-ascorbic acid (Sigma Aldrich, A8960) for 10 days. At day 12 cells were dissociated using TrypLE express and replated as single cells onto fresh Matrigel hESC-Qualified matrix (Corning) plates into neural differentiation medium (as above) supplemented with 20 μM DAPT (Chem Cruz, sc-201315), 10 ng/ml brain-derived neurotrophic factor (BDNF, PeproTech, 450-02-2UG), 10 ng/ml glial-derived neurotrophic factor (GDNF, PeproTech, 450-10-2UG), 1 μM cAMP (Sigma Aldrich, A6885-100mg) and 200 μM L-ascorbic acid (Sigma Aldrich, A8960). Medium was supplemented with 10 μM Y-27632 (Tocris) for the first 24h. During neural induction and maintenance, growth medium was replaced every 48h until day 24.

### Neural crest differentiation

To differentiate NC cells, 80-90 % confluent PNP/NC cultures at P5 were dissociated to single cells and plated at 1:10 onto Matrigel hESC-Qualified matrix (Corning) into DMEM:F12 (Thermo Fisher Scientific) supplemented with 1x B27 supplement (Thermo Fisher Scientific) and 1 % Fetal Bovine Serum (FBS, Sigma Aldrich, F754) (Mohlin et al., 2019). Medium was replenished every 48h for 7 days.

### Trunk-to-tail transition

PNPs were generated and maintained as described above for PNP long term maintenance. Cultures between P25 and P30 were split into long-term PNP maintenance medium (+SBSAG/+SBLDN), NMP differentiation medium (-SBSAG/-SBLDN) or NMP differentiation medium supplemented with 50 ng/ml GDF11 (Peprotech, 120-11-B). Samples were collected for RNA analysis when confluent (48-72h).

### Immunofluorescence microscopy

Cells were cultured in 8 or 12 well μ-slides (Ibidi) and fixed by adding ice-cold 4% Pierce formaldehyde (w/v) methanol-free (Thermo Fisher Scientific, 28908) in PBS for 10-15 mins. Cells were permeabilised in PBS supplemented with 0.1 % (v/v) Triton-X100 (Sigma Aldrich, T8787-250ML) for 10 mins and then blocked solution consisting of PBS supplemented with 0.1 % (v/v) Triton-X100 (Sigma Aldrich), 5% (v/v) Donkey serum (Merck Millipore, S30-100ML) for more than 1h at room temperature. Primary antibodies were incubated in blocking solution at 4°C overnight in concentrations detailed in Supplementary file 6. Cells were then washed in PBS and incubated in Donkey AlexaFluor conjugated secondary antibodies (Abcam) diluted at 1:400 in blocking solution for more than 1 hour at room temperature. Cells were mounted in Vectorshield antifade mounting medium containing DAPI (Vector Laboratories, H-1200-10). Cells were imaged using two imaging systems; 1) by a Zeiss LSM710 confocal microscope (Carl Ziess AG) using Zeiss Plan-Apochromat 20x/0.8 or 10x/0.45 objective (Carl Ziess AG) controlled by ZEN Black 2012 software (Carl Ziess AG); and 2) by an inverted Olympus IX83 microscope (Olympus Corporation) using an Olympus super-apochromatic 20x/0.75 objective (Olympus Corporation), captured using a Hamamatsu Flash 4.0 sCMOS camera (Hamamatsu photonics), a Spectra X(LED) light-source (Lumencore) and controlled by CellSens Dimension software (Olympus Corporation)). Post-acquisition analysis was performed using (Fiji) Image J (Schindelin et al., 2012). Briefly, nuclear segmentation was achieved using a fixed binary threshold using DAPI, the fluorescence intensity (mean grey value) of each channel was masked back to nuclei.

### Flow Cytometry

Cells were collected using Gibco TrypLE express (Thermo Fisher Scientific) dissociation, fixed by adding ice-cold 4% Pierce formaldehyde (w/v) methanol-free (Thermo Fisher Scientific) in PBS for 15 mins, and washed using PBS. Cells were permeabilised with PBS/0.5 % Triton-X100 for 15m and blocked with PBS supplemented with 0.1 % (v/v) Triton-X100 (Sigma Aldrich), 1 % BSA fraction V (w/v) (Sigma-Aldrich, A3059) for 1hr while mixing on a slow speed gyratory motion shaker. Primary incubations were completed in blocking buffer using Alexa Fluor 488 Mouse anti-SOX2 (BD Pharmingen, O30-678) and Alexa Fluor^®^ 647 Mouse anti-CDX-2 (BD Pharmingen, M39-711,). After washes, fluorescence was immediately measured on a LSR II cytometer (BD Biosciences) and results were analysed using FlowJo software (FlowJo LLC). Gates used to determine percentage of positive cells were designed based on fluorescence levels detected in the control samples, which included both Alexa Fluor 488 Mouse IgG1 κ (MOPC-21, BD Pharmingen and Alexa Fluor 647 Mouse IgG1 κ (BD Pharmingen, MOPC-31C) isotype control isotype and unstained sample. Aldehyde dehydrogenase activity was measured as per the manufacturer’s guidelines using the ALDEFLUOR Kit (STEMCELL Technologies, 01700). Fluorescence was measured on a LSR II cytometer (BD Biosciences) and analysed using FlowJo software (FlowJo LLC).

### Clonal expansion of PNPs and NC cells

To generate sub-clonal PNP and NC cell lines, passage 5 cells were selectively detached and dissociated into single cells using TrypLE express (Thermo Fisher Scientific) as previously described. Cells were resuspended into RPMI 1640 (Thermo Fisher Scientific, 32404-014) supplemented with 10% (v/v) KnockOut serum replacement, (KSR, Thermo Fisher Scientific, 10828028) and 10 μM Y-27632 (Tocris). Cells were sorted using a MoFlo XPD (Beckman Coulter) using FSC and SSC profile to select single, live cells. Cells were sorted into Matrigel hESC-Qualified Matrix (Corning) coated 96 well plates (Corning) containing NMP differentiation medium. Surviving cells were subsequently passaged TrypLE express (Thermo Fisher Scientific) to expand clonal population as previously described above.

### RNA extraction, cDNA synthesis and qPCR

Total RNA extraction was completed using RNEasy mini kit (Qiagen, 74106) following the manufacturer’s instructions. cDNA was synthesised using Maxima First Strand cDNA Synthesis Kit for RT-qPCR with dsDNase (Thermo Fisher Scientific, K1672) following manufacturer’s instructions with the addition of a dilution step where cDNA was diluted 1:60 in water. qPCR analysis was performed using primers detailed in Supplementary file 7 on a Roche Lightcycler 480 II (Roche Holding AG) using LightCycler 480 SYBR Green I Master mix (Roche Holding AG, 04887352001). Relative expression was calculated using the ΔΔCt method, normalising each gene to porphobilinogen deaminase (PBGD) levels.

### RNA-sequencing

RNA was extracted using RNEasy mini kit (Qiagen) following the manufacturer’s instructions including recommended DNase digestion step. RNA concentration was measured on a on a GloMax (Promega Corporation) and RNA integrity on TapeStation (Agilent Technologies). Libraries were prepared using KAPA mRNA (PolyA) HyperPrep Kit (Roche Holding AG, KK8581) using 500 ng RNA per sample according to manufacturer’s instructions. Libraries were sequenced using a HiSeq 4000 (Illumina Biotechnology) as follows: pooled to 4 nM, 75bp single end sequencing and up to 38 million reads per sample. Data is available at the GEO repository (accession number GSE150709).

### RNA-seq analysis

Reads were Illumina adapter trimmed using Cutadapt v1.16 (Martin, 2011) and aligned against GRCh38 and Ensembl release 86 transcript annotations using STAR v2.5.2b (Dobin et al., 2013) via the transcript quantification software RSEM v1.3.0 (Li and Dewey, 2011). Gene-level counts were rounded to integers and subsequently used for differential expression analysis with DESeq2 (Love et al., 2014). Differential expression analysis between pairwise replicate groups was thresholded for significance based on an FDR<=0.01, a fold-change of +/− 2, and a base-mean expression of >=100. PCA analysis was conducted on the normalised log transformed count data using the 10% most variable genes across samples. The volcano plot depicts the FDR and logFC statistics from the group DESeq2 differential expression analysis between P5 epithelial and P5 mesenchymal samples. For hierarchical clustering analysis, genes that maintained their significance and direction of change across 2 consecutive time-points were selected for visualisation in a heatmap. K-means clustering (k=10) was used to identify distinct gene clusters of related expression. Heatmaps show gene-level normalised counts, centred and scaled as z-scores. Gene ontology analysis was carried out using ToppGene Suite (ToppFun function) (Chen et al., 2009).

### Comparison between data sets

Previously published Affymetrix array data were downloaded from the NCBI Gene Expression Omnibus (GEO) as GSE109267 (Frith et al., 2018). Cell files were imported into R and RMA processed using the Bioconductor package oligo with default settings. Differential expression analysis between NMP and hESC replicate groups was assessed using limma (Ritchie et al., 2015). Genes with an FDR corrected p-value <= 0.01 and fold change >= +/− 2 were called significant. NMP high genes from the Verrier et al (2018) study were provided in supplementary data and subsequently filtered using a P-value of <=0.01 (Verrier et al., 2018). The overlap between each genes list representing significantly upregulated genes at 36h was generated using BioVenn (Hulsen et al., 2008). The overlap between each gene list was found to be significant (p<1e-4, hypergeometric distribution).

## SUPPLEMENTARY FIGURES

**Figure S1:**
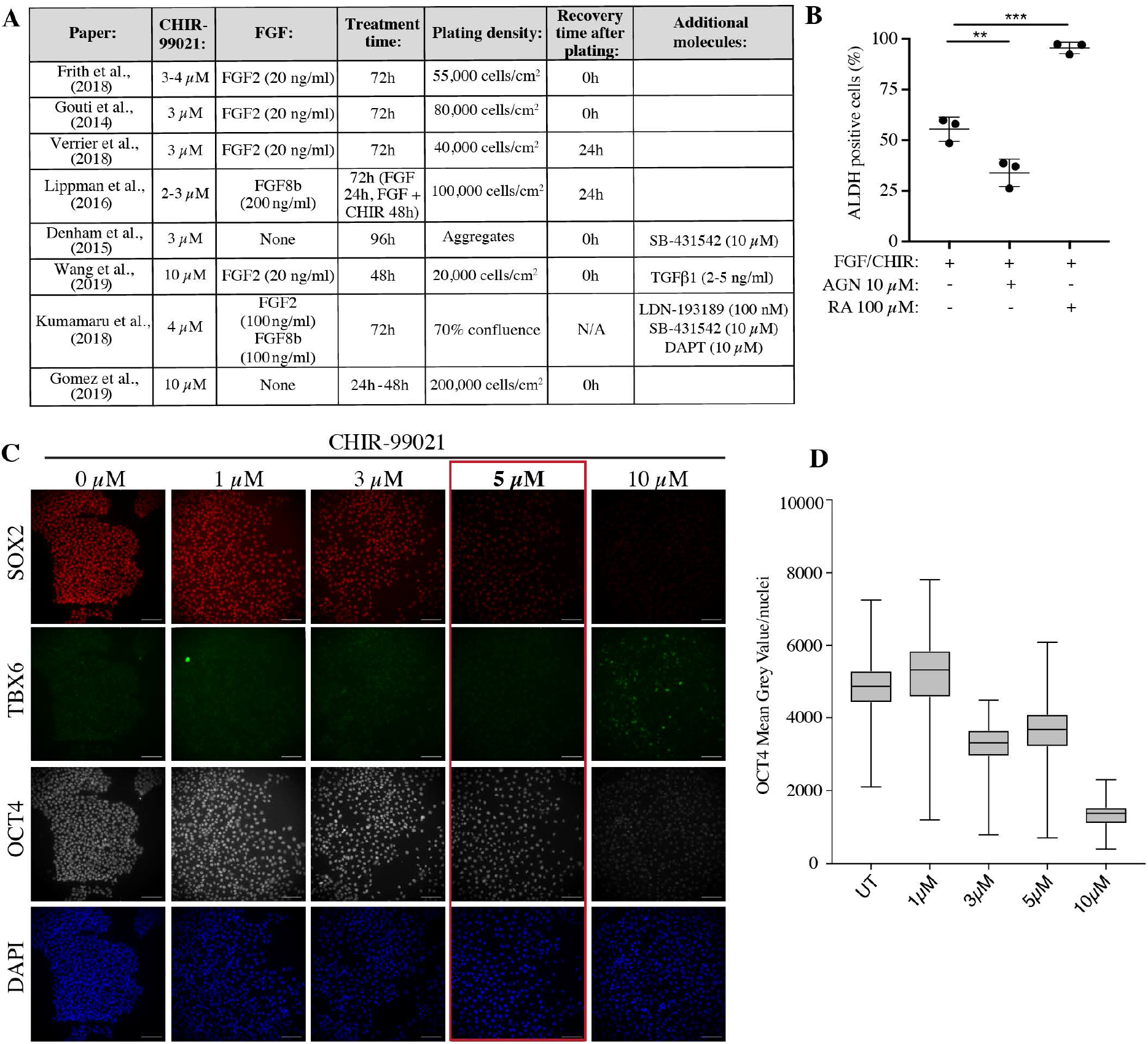
NMP-like cells are induced by combined Wnt/FGF and inhibited RA signalling. A) Summary of protocols used in recent studies to generate NMP-like cells from hPSCs. Table includes plating density and recovery time after plating, as well as the exogenous molecules and treatment time used. B) ALDEFLUOR assay was used to measure the expression of aldehyde dehydrogenases (ALDH) in 36h samples generated in three conditions: 1) FGF2 and CHIR only, 2) FGF, CHIR and AGN or FGF, CHIR and RA. Samples were analysed using flow cytometry and results were presented as the percentage of cells expressing ALDH. Error bars show SD (n = 3 experiments). **P <0.01, ***P <0.001 (ANOVA). C) Representative immunostaining SOX2 (red), TBX6 (green) OCT4 (grey) and the nuclear stain DAPI (blue) after 36h treatment following scheme as shown in Figure 1A with 0 μM, 1 μM, 3 μM, 5 μM and 10 μM CHIR-99021. Scale bars, 100 μm. D) Box-plot showing mean grey value/nuclei quantified from repeat experiments as shown in (C). Plot show data points collected from 2 experiments (>450 nuclei/experiment).

**Figure S2:**
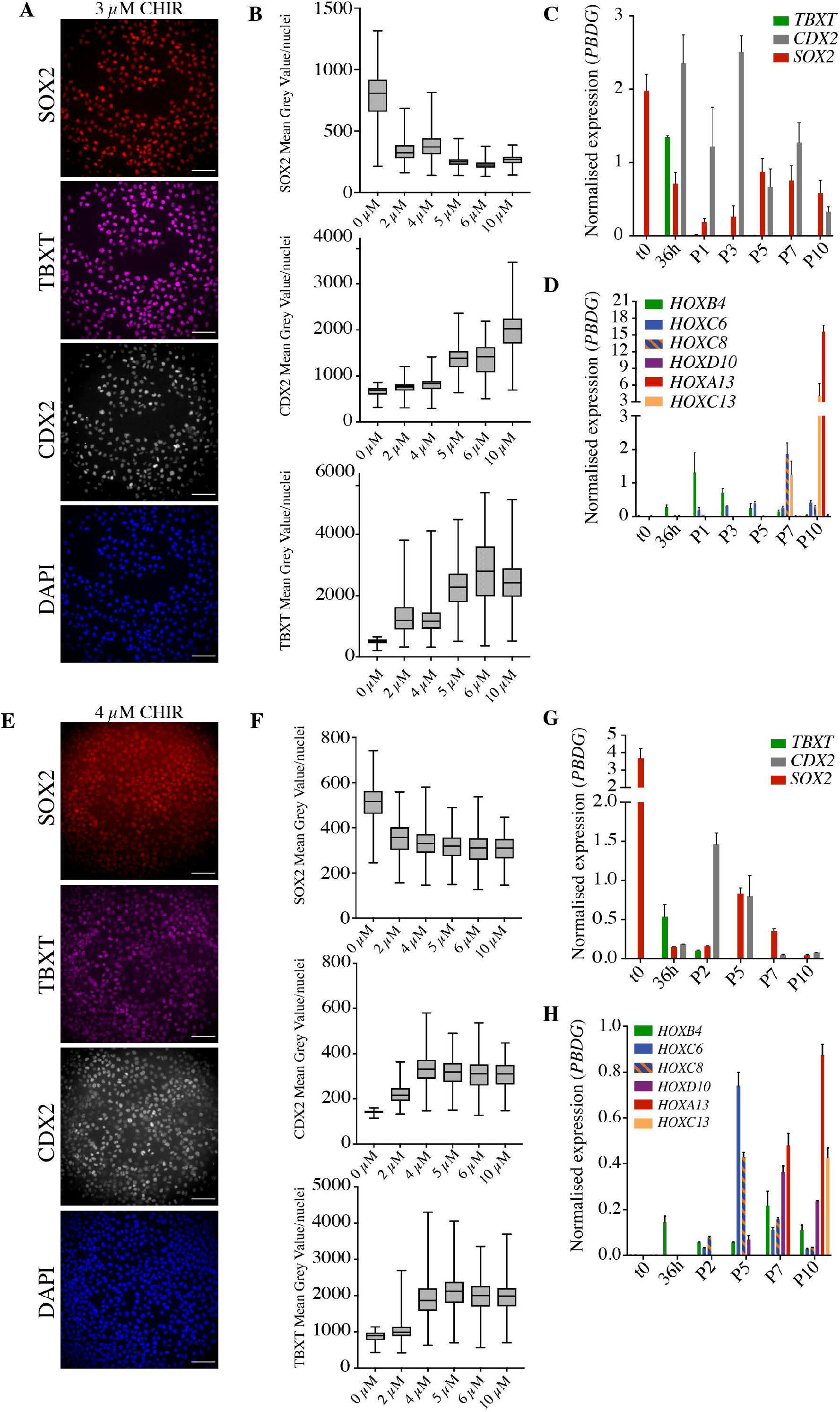
Generation of NMP-like cells in multiple hPSC lines requires modulation of the Wnt pathway. A, B) Optimal CHIR concentration (3 μM) was optimised in WA01 (H1) hESCs. (A) Representative immunostaining of NMP markers SOX2 (red) and CDX2 (grey) and TBXT (magenta) at 36h after following treatment scheme with 3 μM CHIR and (B) quantification markers over a range of CHIR concentrations between 1-10 μM. Scale bars, 100μm. C,D) Transcriptional analysis (RT-qPCR) of NMP markers *TBXT, SOX2* and *CDX2* (C) and selected HOX genes (D) up to passage 10. Expression levels are normalised to the reference gene *PBDG*. Error bars show SD, (n=3 technical replicates). E, F) Optimal CHIR concentration (4 μM) was optimised in AICS ZO1-mEGFP (AICS-0024) iPSCs hESCs. (E) Representative immunostaining of NMP markers SOX2 (red) and CDX2 (grey) and TBXT (magenta) at 36h after following treatment scheme with 3 μM CHIR and (F) quantification markers over a range of CHIR concentrations between 1-10 μM. Scale bars, 100μm. G,H) Transcriptional analysis (RT-qPCR) of NMP markers *TBXT, SOX2* and *CDX2* (G) and selected HOX genes (H) up to passage 10. Expression levels are normalised to the reference gene *PBDG*. Error bars show SD, (n=3 technical replicates).

**Figure S3:**
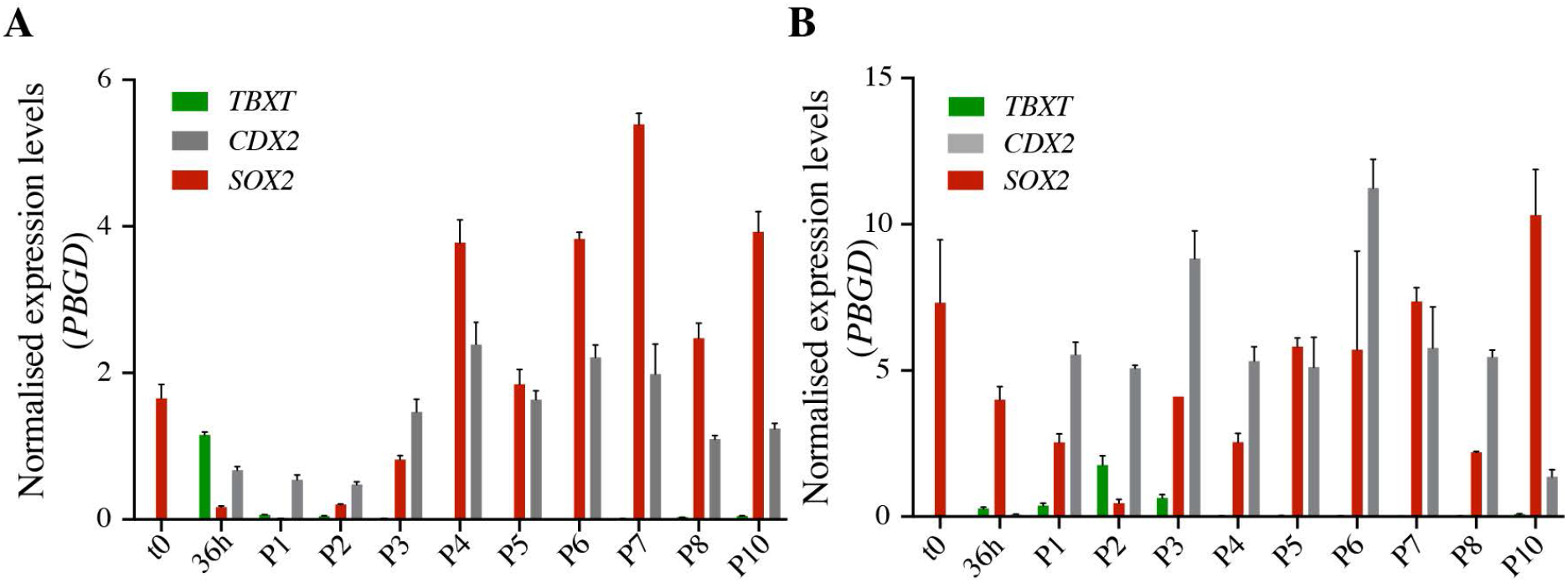
CDX2 and SOX2 expression can be maintained for 10 passages. A,B) Transcriptional analysis (RT-qPCR) of two independent experiments showing NMP markers *TBXT, SOX2* and *CDX2* at each passage, up to passage 10. Expression levels are normalised to the reference gene *PBDG*. Error bars show SD, (n=3 technical replicates).

**Figure S4:**
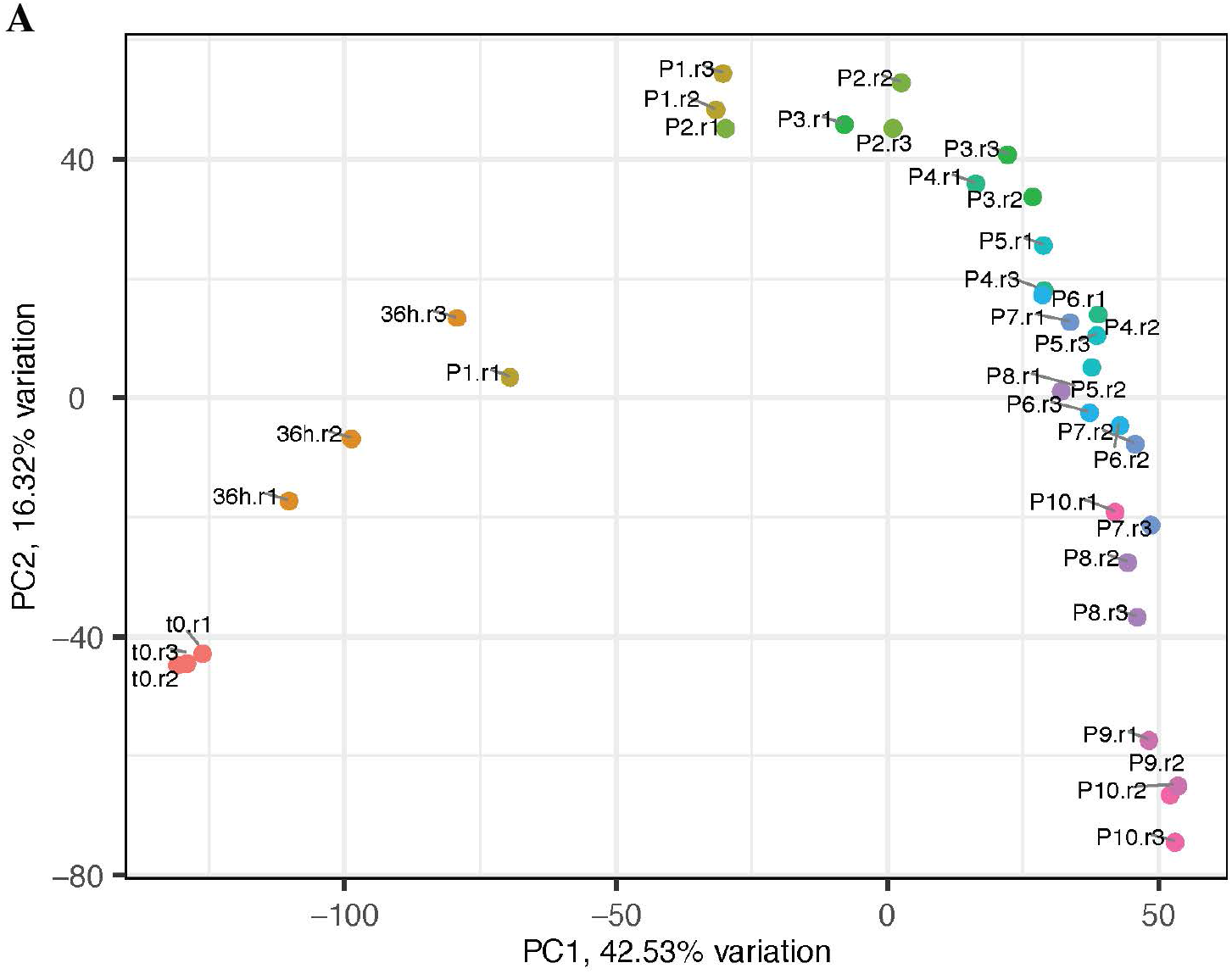
Principal component analysis of RNA-Seq samples collected over passaging. A) PCA analysis show biological replicates for each passage cluster together and show small biological variation between experiments.

**Figure S5:**
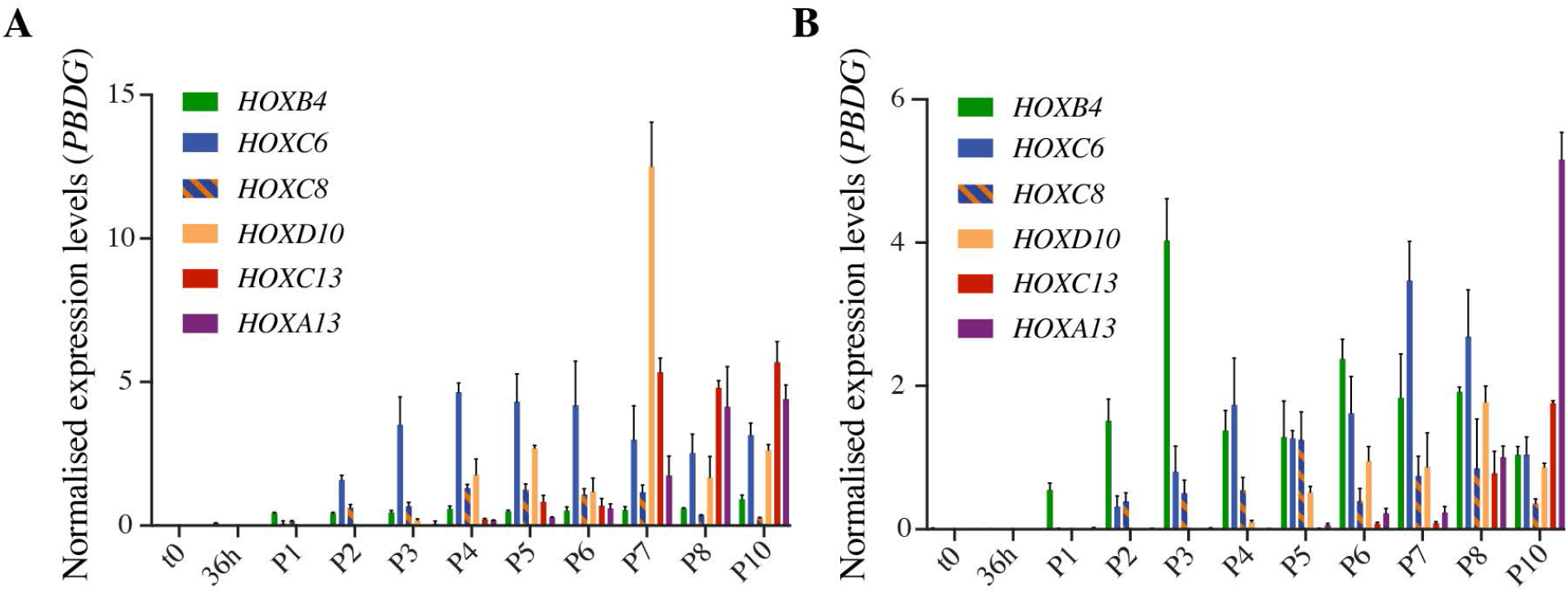
Full collinear expression of the HOX gene cluster occurs over 10 passages. A,B) Transcriptional analysis (RT-qPCR) of two independent experiments showing selected HOX genes at each passage up to passage 10. Expression levels are normalised to the reference gene *PBDG*. Error bars show SD, (n=3 technical replicates).

**Figure S6:**
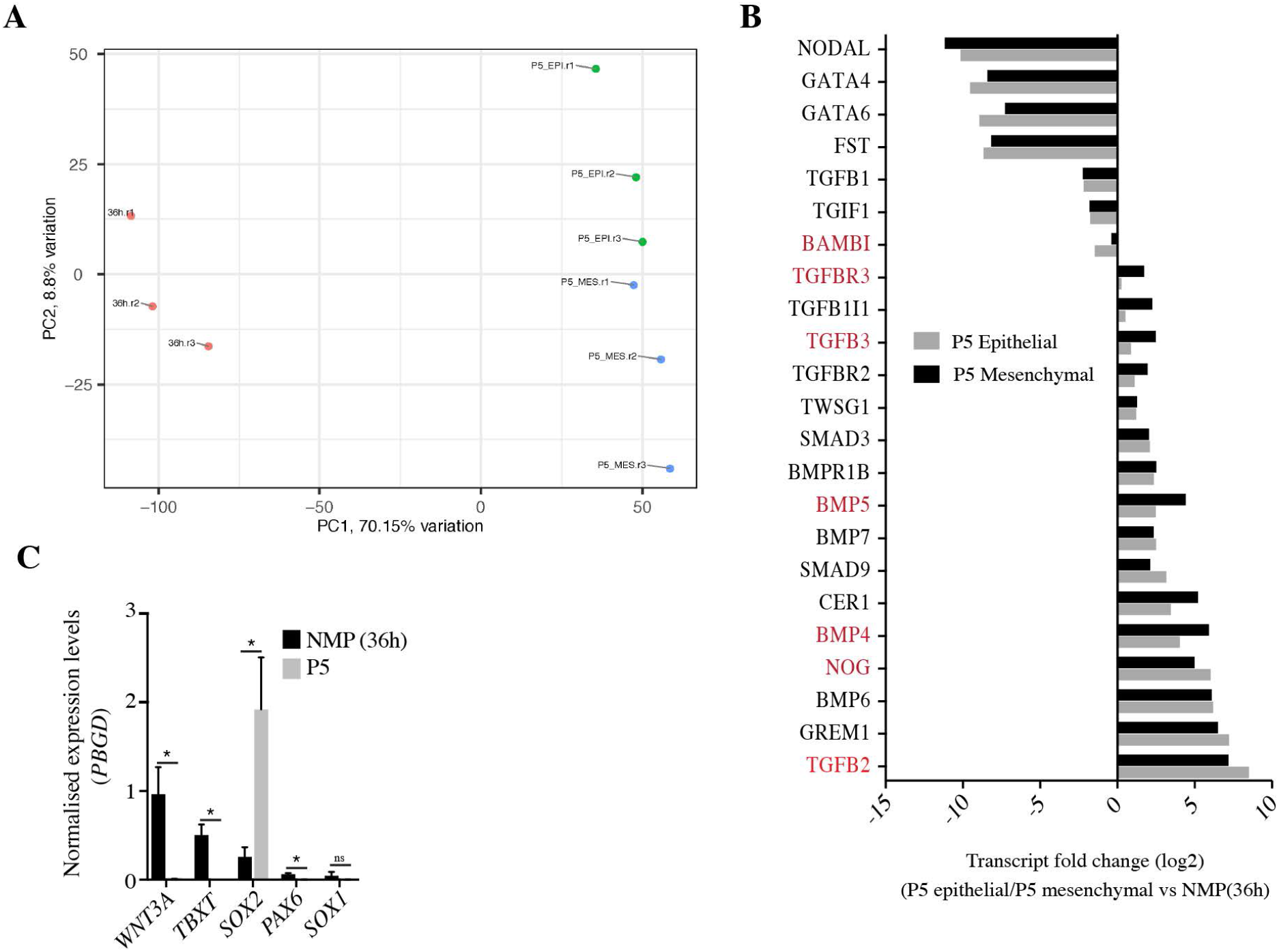
Principal component analysis of mesenchymal and epithelial samples analysed by bulk RNA-sequencing. A) PCA analysis showing biological replicates for the mesenchymal (MES) and epithelial (EPI) enriched samples and NMP samples (36h). B) Graph showing transcriptional fold change (FC) of selected TGF superfamily genes in P5 epithelial and P5 mesenchymal samples over 36h samples. Genes which are statistically differentially expressed between epithelial and mesenchymal samples are highlighted in red. C) Transcript levels of *WNT3A, TBXT, SOX2 PAX6* and *SOX1* in NMP (36h) and P5 samples as measured by RT-qPCR. Expression levels were normalised to the reference gene *PBGD*. Error bars show SEM (n=2/3 experiments), *P <0.05 (unpaired t-test).

**Figure S7:**
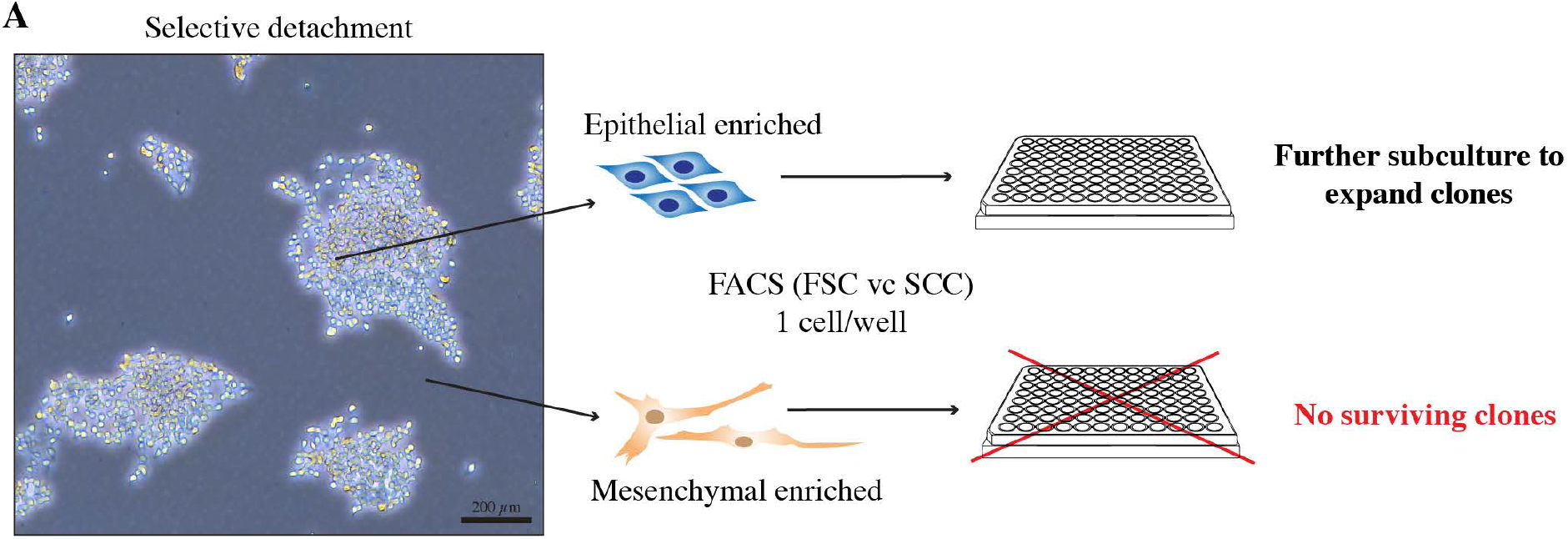
Generating sub-clonal populations from PNP/NC cell enriched samples. A) Scheme to generate sub-clonal populations from mesenchymal- or epithelial-enriched samples. Cells were selectively detached to separate epithelial from mesenchymal cell populations and single cells from each enriched cell sample were sorted (FACS) into wells of a 96 well plate. Surviving sub-clones were expanded for analysis.

**Figure S8:**
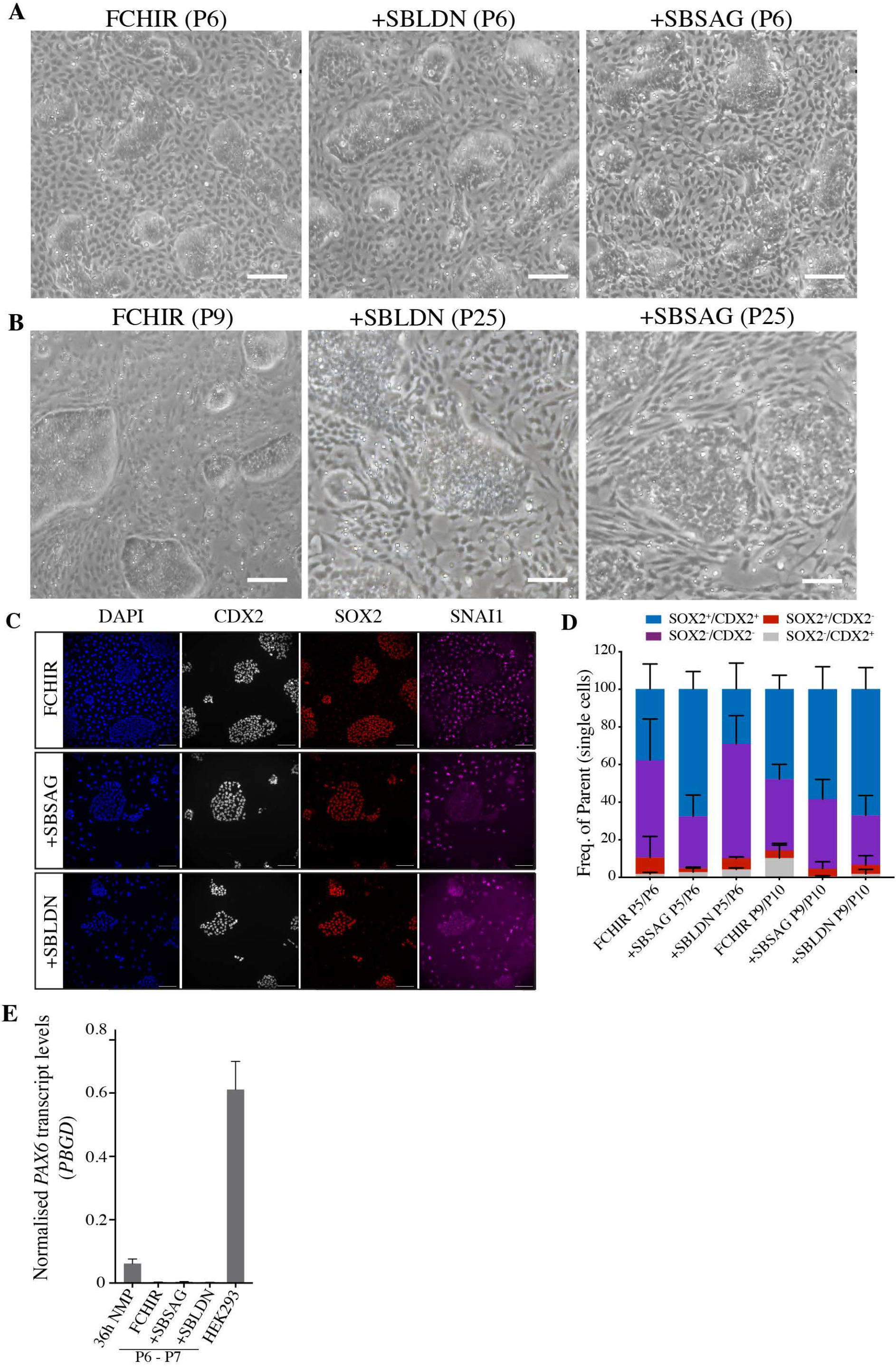
Upregulation of terminal *HOX* genes is significantly delayed, and neural marker gene *PAX6* remains silent, in +SBSAG and +SBLDN conditions. A, B) Representative brightfield images and PNPs/NC at mid (P5) and late passages (FCHIR:P10, +SBLDN and +SBSAG: P25). Scale bar, 200 μm. C) Representative immunostaining of P5 cells for CDX2 (grey), SOX2 (red) and SNAI1 (magenta) under conditions indicated in Figure 7D. Scale bar, 100 μM. D) SOX2/CDX2 flow cytometry analysis of FCHIR and +SBLDN and +SBSAG samples at early and late passages. Cells were analysed using SOX2 and CDX2 conjugated antibodies and plotted as percentage of expression. Error bars show mean with SEM (n = 3). C) Quantification of *PAX6* transcript levels under various conditions as indicated in Figure 7D, and in comparison to HEK293 (positive control) cells. Expression levels were normalised to reference gene *PBGD.* Error bars show mean with SEM (n = 2-3).

**Figure S9:**
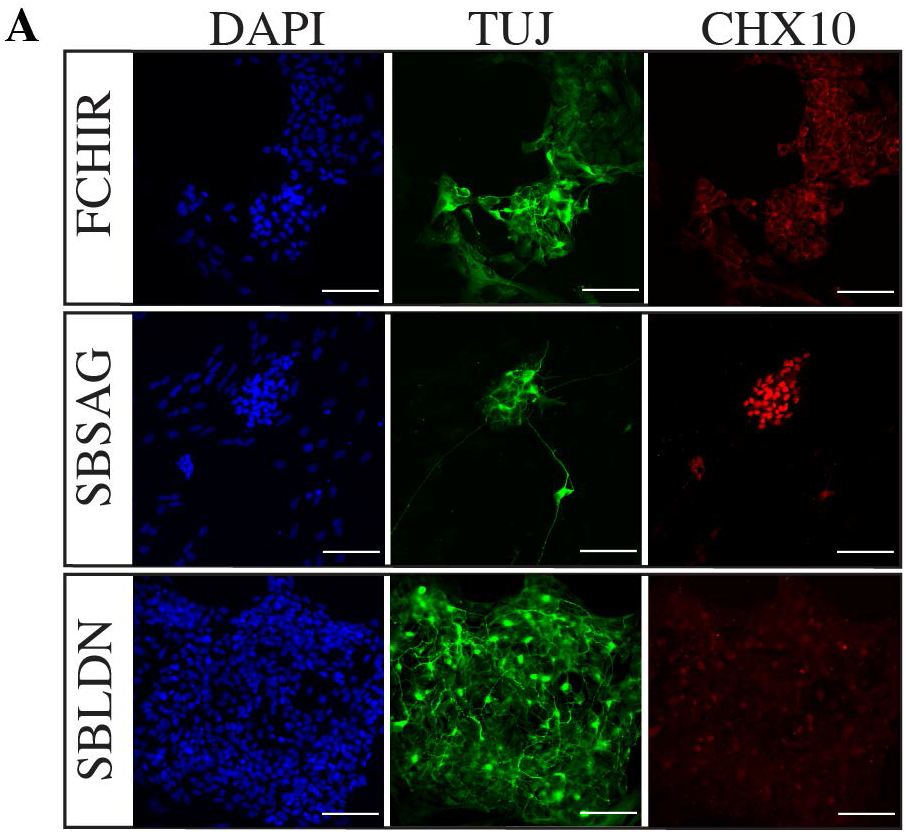
NMP-derived PNPs treated with SHH generate ventralised neuronal cultures. A) Representative immunostaining of ventral neurons stained with CHX10 (red) paired with βIII-tubulin (TUJ, green) and nuclear stain DAPI (blue). Scale bars, 100μm.

